# Origin and cross-century dynamics of an avian hybrid zone

**DOI:** 10.1101/012856

**Authors:** Andrea Morales-Rozo, Elkin A. Tenorio, Matthew D. Carling, Carlos Daniel Cadena

**Affiliations:** Laboratorio de Biología Evolutiva de Vertebrados, Departamento de Ciencias Biológicas, Universidad de Los Andes, Bogotá, Colombia; Calima: Fundación para la Investigación de la Biodiversidad y Conservación en el Trópico, Cali, Colombia.; Cornell Laboratory of Ornithology, Cornell University, Ithaca, NY, USA; Department of Zoology and Physiology, University of Wyoming, Laramie, WY, USA; Programa de Biología y Museo de Historia Natural, Universidad de los Llanos, Sede Barcelona, Villavicencio, Colombia

**Keywords:** Andes, cline, Hill function, distribution modeling, hybridization, moving hybrid zone

## Abstract

**Background:** Characterizations of the dynamics of hybrid zones in space and time can give insights about traits and processes important in population divergence and speciation. We characterized a hybrid zone between tanagers in the genus *Ramphocelus* (Aves, Thraupidae) located in southwestern Colombia. We tested whether this hybrid zone originated as a result of secondary contact or of primary differentiation, and described its dynamics across time using spatial analyses of molecular, morphological, and coloration data in combination with paleodistribution modeling.

**Results:** Models of potential historical distributions based on climatic data and genetic signatures of demographic expansion suggested that the hybrid zone likely originated following secondary contact between populations that expanded their ranges out of isolated areas in the Quaternary. Concordant patterns of variation in phenotypic characters across the hybrid zone and its narrow extent are suggestive of a tension zone, maintained by a balance between dispersal and selection against hybrids. Estimates of phenotypic cline parameters obtained using specimens collected over nearly a century revealed that, in recent decades, the zone appears to have moved to the east and to higher elevations, and has apparently become narrower. Genetic variation was not clearly structured along the hybrid zone, but comparisons between historical and contemporary specimens suggested that temporal changes in its genetic makeup may also have occurred.

**Conclusions:** Our data suggest that the hybrid zone likey resulted from secondary contact between populations. The observed changes in the hybrid zone may be a result of sexual selection, asymmetric gene flow, or environmental change.

## Background

Characterizations of hybrid zones allow one to make inferences about traits and processes relevant to understanding the origin and maintenance of differences between populations and species [1,2]. A classic question about hybrid zones is how are they formed, with previous studies proposing two main hypotheses (reviewed by [3]). The hypothesis of secondary contact posits that hybrid zones result from expansion of populations that were previously isolated geographically and which interbreed in contact zones because complete reproductive isolation between them was not reached during the allopatric phase [4]. An alternative hypothesis postulates that hybrid zones form in parapatry, by primary differentiation across ecological gradients [5]. Secondary contact is likely if environments that presently allow the distributions of hybridizing populations to overlap were disjunct in the past, a scenario that predicts one should observe genetic signatures of population expansions. Alternatively, primary differentiation along a gradient would occur if the extent of suitable environments for the hybridizing populations has been stable over time; this predicts that populations have not expanded their ranges historically, and that the position of the hybrid zone (as indicated by the position of clines in molecular and morphological traits) is coincident with an environmental transition [1,6].

Inferring hybrid-zone origins from current patterns of variation is challenging because, with time, genetic signatures of secondary contact or primary intergradation tend to erode [3,7]. Alternatively, then, tests of hypotheses posed to account for the origin of hybrid zones may be conducted by examining the historical distribution of hybridizing taxa using paleodistributional modeling, i.e. projecting ecological niche models, which characterize the current distribution of species in climatic space, onto models of historical climatic conditions to infer potential distributions in the past [8,9]. Such models of historical distributions represent hypotheses one can further test using molecular data to evaluate their population-genetic predictions, such as signatures of population growth for presumably expanding populations and of constant population size and isolation by distance in populations occurring within climatically stable areas [10,11]. This approach has revealed that several hybrid zones likely originated following range expansions leading to secondary contact [12-15].

Another focus of studies on hybrid zones is the analysis of their temporal dynamics, which can allow understanding the role played by different evolutionary forces in such scenarios. When hybrid genotypes are less fit than parental genotypes, ‘tension zones’ are formed, which are maintained by a balance between the homogenizing effect of dispersal into the hybrid zone and the diversifying effect of selection against hybrids [1]. If there is endogenous selection against hybrids, then there should be coincidence in location and concordance in width of clines describing the variation in different traits and loci across a hybrid zone, and such clines should remain stable over time [16,17]. However, hybrid zones are often temporally dynamic (i.e. they may shift in location or change in width) and because clines for different traits may change in different ways, one can make inferences about the action of particular processes (e.g., natural selection, sexual selection, competition, asymmetric hybridization, dominance drive) based on dynamics observed for different characters [18]. For example, discordant patterns of plumage and mitochondrial DNA variation across a hybrid zone between *Setophaga* warblers, coupled with behavioral experiments showing aggressive superiority of males of one species over the other, indicate that movement of this zone has likely been driven by competition-mediated asymmetric hybridization [19-21]. Temporal changes in the makeup of hybrid zones may also reflect natural or human-mediated environmental changes [22,23].

There is ample evidence of hybridization between members of the tanager genus *Ramphocelus* (Aves, Thraupidae; [24-27]), but detailed studies on hybrid zones involving species in this group are scant. Here, we characterize a hybrid zone between members of this genus located in western Colombia that has received little study although its existence was noted nearly a century ago [28] and was described in some detail more than five decades ago [25]. In the Cauca River Valley above c. 900 m elevation, one finds the larger form *flammigerus* (males are black with a scarlet rump), whereas the smaller and yellow-rumped form *icteronotus* occurs along the costal plains west of the Andes extending north into Costa Rica and south into northern Peru. Females and immatures are similar to their respective males, but are less strongly colored. Along a c. 140 km transect running approximately northwest from the city of Cali downslope along the western flank of the Cordillera Occidental, the two forms hybridize, forming a gradient in coloration and body mass ([25,29]; Fig. 1 and 2). Currently, these forms are considered subspecies of *Ramphocelus flammigerus* [30] because gene exchange between them appears to be unrestricted, with apparently no selection against hybrids [25].

**Fig. 1.**
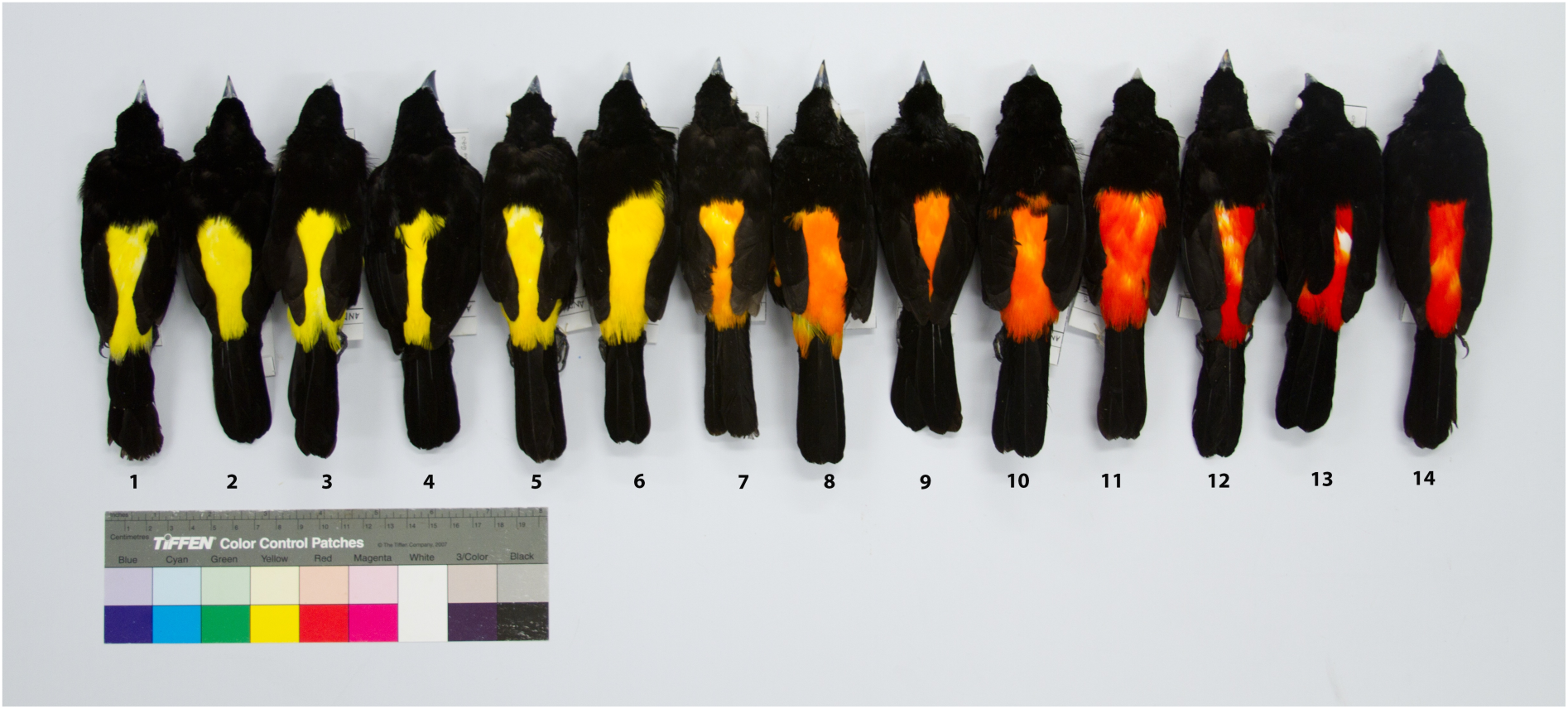
Phenotypic variation in male specimens collected along the *Ramphocelus flammigerus* hybrid zone in southwestern Colombia. Individuals 1-6 correspond to *R. flammigerus icteronotus* (yellow-rumped form) from the plains of the Pacific coast (sector 1; see Fig. 2). On the other extreme, individuals 11-14 correspond to *R. flammigerus flammigerus* (scarlet-rumped form) distributed towards the Cauca River Valley (sector 3). Individuals 7-10 are intermediates collected near the center of the hybrid zone (sector 2).

**Fig. 2.**
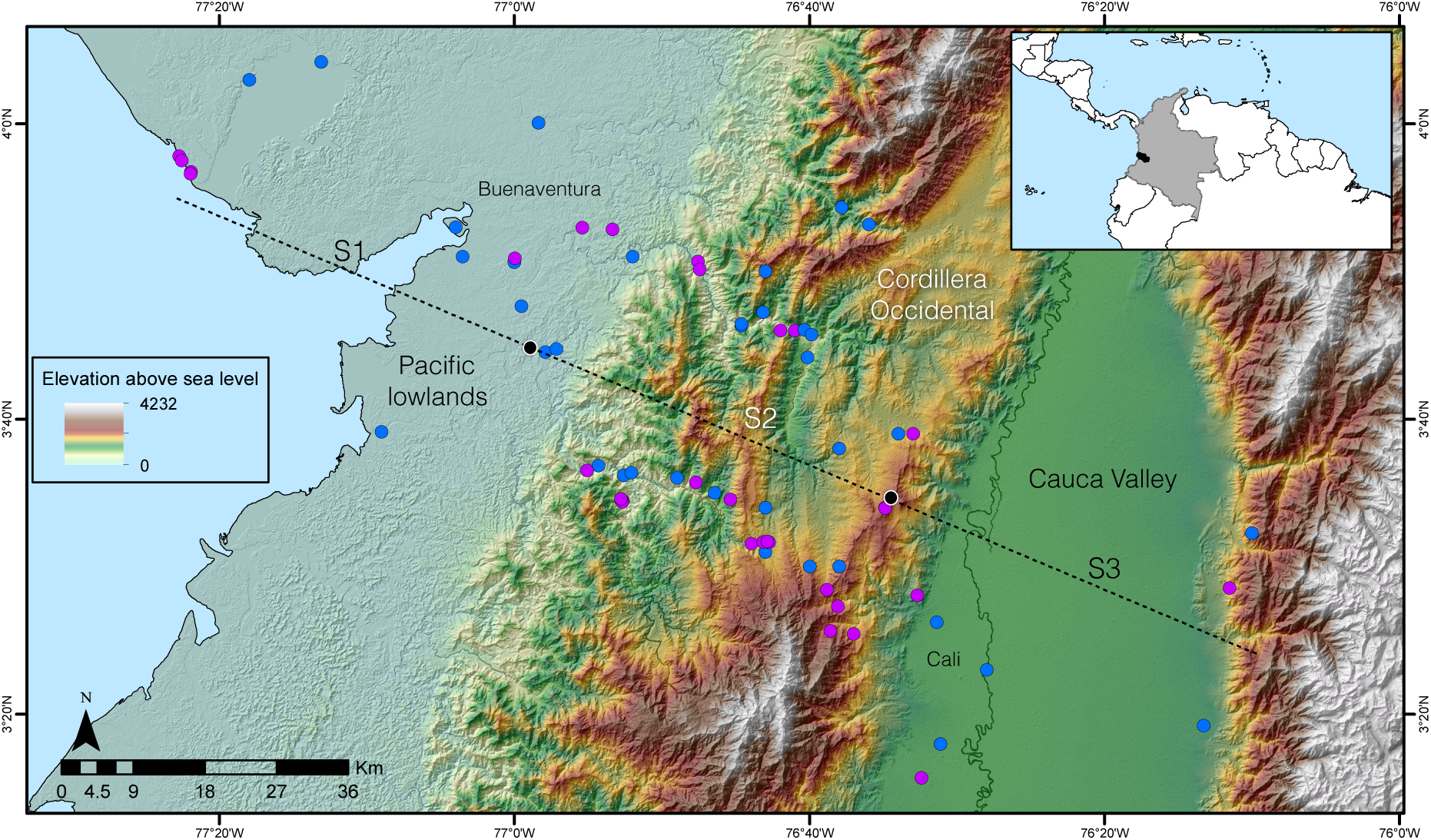
Study transect encompassing the *Ramphocelus* hybrid zone from the Pacific lowlands to the eastern slope of the Cordillera Occidental of the Colombian Andes. Blue dots indicate collection sites of historical specimens and purple dots indicate collection sites of current specimens sampled for phenotypic/genetic variation; black lines correspond to municipality limits, with text indicating the location of the larger cities of Cali and Buenaventura. The extent of each of the three sectors we defined in the hybrid zone (S1, S2 and S3; see text) is indicated by black dots on the transect.

The *R. flammigerus* system is particularly well suited to studying the role of different evolutionary forces at work in hybrid zones owing to the existence of variation in characters with different modes of inheritance, and, presumably, under different forms of selection. Variation in rump coloration across the hybrid zone likely reflects variation in the concentration of a single carotenoid pigment and is influenced by the environment because carotenoids are obtained from the diet [25,31,32]. Furthermore, based on the strong sexual dichromatism in this species, and the likelihood that its mating system involves some degree of polygamy [33], plumage coloration is probably influenced by sexual selection. In contrast, morphometric variation [25,29] likely has a strong genetic basis [34-36] and could be subject to natural selection [37-39]. The value of considering traits with different modes of inheritance and under different selective pressures to understand evolutionary forces at work in hybrid zones is illustrated by studies showing (1) that clines for traits involved in courtship are displaced with respect to clines for presumably neutral traits or loci, suggesting a role for sexual selection driving introgression [40,41]; (2) that sex-linked molecular markers introgress over shorter distances than autosomal markers, suggesting a role for sex chromosomes in reproductive isolation [42,43]; or (3) that there is more limited introgression in organellar DNA than in nuclear genes, suggesting selection acts more strongly on hybrids of the heterogametic sex (Haldane's rule; [44]).

Here, we sought to evaluate whether the *Ramphocelus* hybrid zone in southwestern Colombia originated as a result of secondary contact or of primary differentiation, and to examine the zone's dynamics over nearly a century to make inferences about the action of different evolutionary processes. To accomplish this, we (1) reconstructed the biogeographic and demographic history of the hybridizing populations based on ecological niche modeling and coalescent analyses of mtDNA sequence data, (2) characterized genetic, morphological and plumage variation across the hybrid zone, and (3) compared spatial patterns of variation in morphometrics, plumage coloration, and genetic structure between specimens collected at different times to assess changes in the position and width of the hybrid zone.

## Materials and methods

### Samples

We characterized the *Ramphocelus* hybrid zone historically by examining specimens collected from 1894 to 1986 in the ornithological collections of the American Museum of Natural History, the Cornell University Museum of Vertebrates, Universidad del Valle, and the Instituto de Ciencias Naturales at Universidad Nacional de Colombia. To describe current patterns of variation, we collected 73 new specimens in 2007-2010: 65 of them are from localities ranging across the hybrid zone over a distance of 140 km by road connecting the cities of Cali and Buenaventura in department Valle del Cauca [25]; the remaining eight are from localities distant from the hybrid zone in the departments of Antioquia (3), Risaralda (3), and Cauca (2). Study skins and tissue samples are deposited in the Museo de Historia Natural de la Universidad de los Andes (ANDES, Table S1). All specimen localities (historical and current) were plotted and their position along a transect line that best adjusted to points (estimated using a linear regression between latitude and longitude) was recorded. To construct character clines, we recorded the perpendicular position of each specimen on the regression line (Fig. 2; Table S1) and calculated the distance from the northwest extreme of our study transect on the Pacific coast to the position of each specimen along the line.

### Biogeographic history

To model potential distributions of the study taxa, we used 343 georeferenced localities obtained from museum specimens and reliable field observations from Costa Rica, Panama, Colombia, Ecuador, and Peru ([45]; GBIF Data Portal, C. Sánchez, pers. comm., our observations). We associated localities with GIS layers for 19 climate variables at a c. 1 km resolution developed for the present (WorldClim; [46]). With these data, we used a maximum entropy approach (Maxent; [47]) based on current climate layers to generate models of the ecological niche and potential distribution of *flammigerus* and *icteronotus* at present (localities of both forms and presumed hybrids were considered together). Model performance in predicting present distributions (evaluated using receiver-operating-characteristic curves; [48]) was satisfactory (see below), validating the use of this approach to infer potential distributions in the past.

We projected models based on current climate data onto historical climate surfaces for 6,000 years ago and for the Last Glacial Maximum (LGM; 21,000 years ago) to determine whether the distributions of our study taxa were likely disjunct in the past as predicted by the secondary contact hypothesis, or have likely been continuous as predicted by the primary intergradation hypothesis. This approach requires assuming ecological niche conservatism and that climate represents a long-term stable constraint on potential distributions. Because ecological niche models are based only on climatic data, they tend to overpredict potential distributions into areas where the study species do not occur owing to historical limitations to dispersal (e.g., the Amazon region in *R. flammigerus*). To reduce overprediction, we cropped maps of current and historical distributions to include only areas within the Andes Ecoregion [49]. Although cropping potential distributions to this region probably did not remove all areas of model overprediction, it allowed for a semi-quantitative comparison of potential distributions across different time periods by calculating the extent of presence areas within the ecoregion. Maxent produces a continuous output ranging from zero to one describing the probability of the species potentially being present at different sites. We considered a threshold of 10% omission to categorize pixels as suitable or unsuitable for each of the time periods.

### Genetic characterization and demographic history

We analyzed variation in DNA sequences of the cytochrome *b* mitochondrial gene for 58 of the *flammigerus/icteronotus* individuals collected in Colombia from 2007 to 2010. In addition, we obtained sequences for three individuals from Ecuador and two from Panama, and combined our data with two sequences of *flammigerus* available in GenBank: one from Ecuador (accession U15719.1; [50]) and one from Panamá (FJ799882.1; [51]; Table S1).

DNA was extracted from tissue samples or toepad samples taken from specimens using a Qiagen DNeasy Tissue Kit or a phenol-chloroform method [52]. PCRs used primers H16064 and L14996 [53] in 24 μl amplification reactions using the following conditions: 42ng of DNA, 0.416 mM dNTPs, 0.5mM of each primer, 1.042 units of 10X buffer with 1.56 mM MgCl2, 0.0246 units/ml AmpliTaq DNA polymerase), and 16.5 of sterile ddH2O. Reactions began with an initial denaturation at 94°C for two minutes, followed by 34 cycles of denaturation at 94°C for 30 s, annealing at 52°C for 30 s, and extension at 72°C for one minute, with a final extension phase at 72°C for 7 minutes. PCR products were purified with Affymetrix Exosap-IT and sequenced in both directions. Sequences were edited, assembled and aligned using Geneious Pro 3.6.1 (www.geneious.com). The mean length of these sequences was 988.8 bp (range 888-1008 bp).

We also analyzed mtDNA from historical toepad samples of 87 specimens collected by C. G. Sibley in 1956 and housed at the Cornell University Museum of Vertebrates [25]. DNA extraction and amplification of these samples was carried out in a historical DNA lab, following protocols to reduce the odds of contamination [54]. For these specimens, we amplified and sequenced c. 210 bp of the cytochrome *b* gene (mean=209.9, range=203-210 bp).

We examined genealogical relationships among haplotypes observed in *flammigerus* and *icteronotus* at present using a maximum-likelihood (ML) phylogenetic analysis employing the GAMMA model. Nodal support was estimated using 1000 bootstrap replicates in RAxML, run from the RAxML BlackBox Web-Server [55]. We used as outgroups sequences of *Ramphocelus carbo* (AF310048.1; [56]) and *R. passerinii* (EF529965.1; [51]).

To assess potential changes in the genetic makeup of the hybrid zone over time, we examined population structure separately for the 1956 specimens and for our samples collected in 2007-2010 (hereafter 2010 specimens) employing procedures implemented in the program ARLEQUIN v3.5 [57]. Based on each data set, we conducted analyses of molecular variance (AMOVAs). We divided our sampling transect in three sectors of equal length: sector 1, 0 – 44 km; sector 2, 45-89 km; and sector 3, 90-134 km. Within each sector, we grouped individuals collected within 1km from each other in a single locality. We calculated F-statistics to estimate differentiation among sectors (F_CT_), among localities within sectors (F_SC_), and among localities among sectors (F_ST_). This analysis allowed us to examine whether there has been any change in the way in which genetic variation is distributed within and among sectors; if spatial genetic structure has become eroded (e.g., if introgressive hybridization has led to genetic homogenization across the transect), then one would expect an increase in genetic variation existing within sectors and a decrease in that existing among sectors over time (i.e. higher F_CT_ in the past than at present). To make data comparable across time periods, sequences for 2010 specimens were trimmed to match the 210 bp available for the 1956 specimens. As an additional way to visualize potential changes in genetic structure over time, we constructed median-joining haplotype networks for the 1956 and 2010 samples using the program PopART (http://popart.otago.ac.nz/index.shtml).

To determine whether populations of *flammigerus* and *icteronotus* have experienced demographic expansions or if these taxa have exhibited historically stable population sizes, we examined trends in effective population size through time using the Extended Bayesian Skyline Plot (EBSP) method implemented in Beast v1.7.4 [58] using sequence data for the 2010 specimens. Demographic expansions are expected if the hybrid zone originated following secondary contact and stable population sizes are expected if the zone originated by primary differentiation. Analyses used the HKY+Γ model, which was selected as the best fit to the data according to the Bayesian Information Criterion (BIC) in JModelTest v 2.1.3 [59]. We ran the analysis for 25,000,000 iterations of which the first 10% were discarded as burn-in; genealogies and model parameters were sampled every 10,000 iterations. For time calibration we assumed a lognormal relaxed clock and a cytochrome−*b* substitution rate of 2.08% divergence per million years [60]. We used the mean of the distribution of population size as a prior (parameter “demographic.populationMean”) calculated from a “Coalescent: constant time” tree prior, run with the same parameters as above. Because this analysis assumes no genetic structure within the sample, we only considered populations located between Cali and Buenaventura (i.e., from the hybrid-zone transect). The skyline plot was built in R [61] with code written by Valderrama *et al*. [62].

### Phenotypic characterization

To characterize the hybrid zone phenotypically, we measured six morphological characters on museum specimens (*n*=139 males, 83 females) with dial calipers to the nearest 0.1 mm: wing length (chord of unflattened wing from bend of wing to longest primary), exposed culmen, bill depth (at the base), bill width (at the base), tail length (from point of insertion of central rectrices to tip of longest rectrix), and tarsus length (from the joint of tarsometatarsus and tibiotarsus to the lateral edge of last undivided scute). To describe morphological variation, we reduced variation in these characters using a principal components analysis (PCA).

We characterized plumage coloration based on reflectance spectra from 400 to 700 nm measured on the rump of adult museum specimens (*n*=144 males, 70 females) using an Ocean Optics USB4F00243 Spectrometer with the SpectraSuite software (Ocean Optics). Three color measurements were estimated for each reflectance spectrum based on segment classification analysis [63]: brightness, an index of how much light is reflected from the sample relative to a white standard; chroma, the saturation of color; and hue, which relates to the wavelength of maximum slope. These measurements were calculated using R code written by Parra [64].

### Phenotypic clines and temporal dynamics

To compare patterns of morphometric and plumage color variation among adult specimens collected at different times in a geographical context, we defined three periods based on temporal sampling gaps: prior to 1911, 1956-1986, and 2010. Hybrid zones are often studied using cline-fitting algorithms that employ Bayesian or maximum-likelihood methods to estimate parameters like cline center and width (e.g. HZAR [65], Analyse [66], ClineFit [67]), and are based on population genetic models. These approaches typically require data from multiple individuals per sampling site to properly characterize variation within and among localities; accordingly, individuals need to be sampled at (or assigned to) a set of discrete sites. This approach is possible when large numbers of specimens are available and when sampling schemes have been explicitly designed with the goal of characterizing variation in space. In our case, many historical specimens were not collected with the specific purpose of describing the *Ramphocelus* hybrid zone and were not sampled at the same set of sites across different time periods. Instead, specimens were collected largely opportunistically at multiple localties and were widely scattered across the study region, with sampling varying spatially and in terms of number of individuals over time. Therefore, because our data did not readily allow us to assign individuals to discrete sampling sites, we did not employ cline-fitting methods; instead, we described the variation in morphometrics and plumage reflectance (i.e. chroma) across the hybrid zone in different time periods using log-logistic models, commonly known as Hill functions [68]. These functions are based on a sigmoidal dose-response (variable slope) model, which describes a response variable y (i.e. character value in our study) as a function of an independent variable x (i.e. distance from the initial point of the hybrid zone) based on a four-parameter logistic equation:

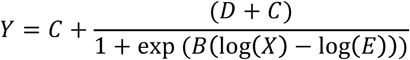

Here, *D* and *C* are the Y values at the plateau’s extremes (i.e. character values at each end of the hybrid zone in our case), *B* is a coefficient denoting the steepness of the curve, and *E* corresponds to the distance along the *X* axis where 50% of the value in *Y* is observed (also denoted ED50; [68]). We used the ED50 value as an estimate of cline center, and estimated cline width as the difference between the ED10 and ED90 values. We used this criterion to estimate cline width because ED10 and ED90 define the values of phenotypes in the *X* axis beyond which there are no intermediate individuals in morphology and color. For example, only individuals with scarlet-rump would be observed in distances in *X* above ED90, whereas only yellow-rumped individuals would be observed in distances below ED10. We calculated the above parameters using the *drm* and *ll.4* functions implemented in the drc package for R [69].

Due to differences in sampling effort over the hybrid zone across time periods, we evaluated the sensitivity of our estimates of cline center and width to sampling effects and calculated confidence limits for these parameters based on a bootstrapping procedure. For each time period and for both morphology and plumage reflectance data, we generated a bootstrap distribution of cline center and width estimated from 1,000 data sets constructed by keeping sample size constant while resampling individuals with replacement. Because in some cases sample size was small, bootstrapping resulted in some samples in which variation was not clinal or in which estimates of cline parameters were unrealistic given the geographic extent of the hybrid zone; therefore, we only retained bootstrap samples in which the estimated cline center took values between 0 and 140 km. We compared estimates of cline center and width across the tree time periods using analyses of variance (ANOVA) and post-hoc Tukey tests treating estimates obtained in bootstrap samples as replicates. Likely due to a low number of individuals with morphological data for the western extreme of the hybrid zone in 2010, bootstrap estimates of cline width for this time period were highly variable and unrealistic; therefore, we do not report confidence limits for this parameter and did not include it in the ANOVA.

### Validation of Hill-function methods

To validate the use of Hill functions to describe phenotypic and genetic clines and to infer cline center and width parameters, we compared our estimates based on Hill functions to estimates obtained by widely used cline-fitting algorithms (HZAR and Analyze) on a published data set that has been subject to cline analyses, namely the molecular data of seven loci of the *Manacus* hybrid zone in Panama [40]. We used the published allele frequency data [40] to estimate cline centers and widths using our approach and compared such estimates to published parameters [65]. Consistent estimates of parameters across methods would validate our Hill-function approach as an alternative method to estimate hybrid zone cline in scenarios where traditional algorithms cannot be used.

## Results

### Biogeographic history

A potential distribution model developed under current climatic conditions in Maxent accurately predicted the present-day distributions of *flammigerus+icteronotus*, with an area under the ROC-curve score of 0.987. Because this suggests that the assumption that climate limits distributions in these taxa is reasonable, we projected models onto past climatic conditions to estimate the potential historical distributions of *flammigerus+icteronotus* at 6,000 and 21,000 years ago.

Although our models evidently overpredict potential distributions, it is clear that the extent of suitable environments for *flammigerus+icteronotus* has not been stable over time. The modeled potential range of these taxa at present in the Northern Andes Ecoregion extends for c. 420,000 km^2^. The predicted potential distribution based on climate for 6,000 years ago was of similar size, with c. 460,000 km^2^ (Fig. 3a). This indicates that current climatic conditions and those from 6,000 years ago were similarly suitable for the presence of these taxa across the study region. Indeed, models suggest that the two forms could have been in contact at that time in the current location of the hybrid zone (Fig. 3a). In contrast, the predicted range during the LGM was considerably smaller than the predicted current range (c. 280,000 km^2^; Fig. 3b). Moreover, suitable conditions for *flammigerus+icteronotus* 21,000 years before present were not continuous along the Pacific slope of the Cordillera Occidental, suggesting that these two forms were likely disjunct during the LGM. Thus, the hybrid zone may have originated following population expansions and secondary contact as a result of climatic change since the LGM, a possibility we address below based on patterns of genetic variation.

**Fig. 3.**
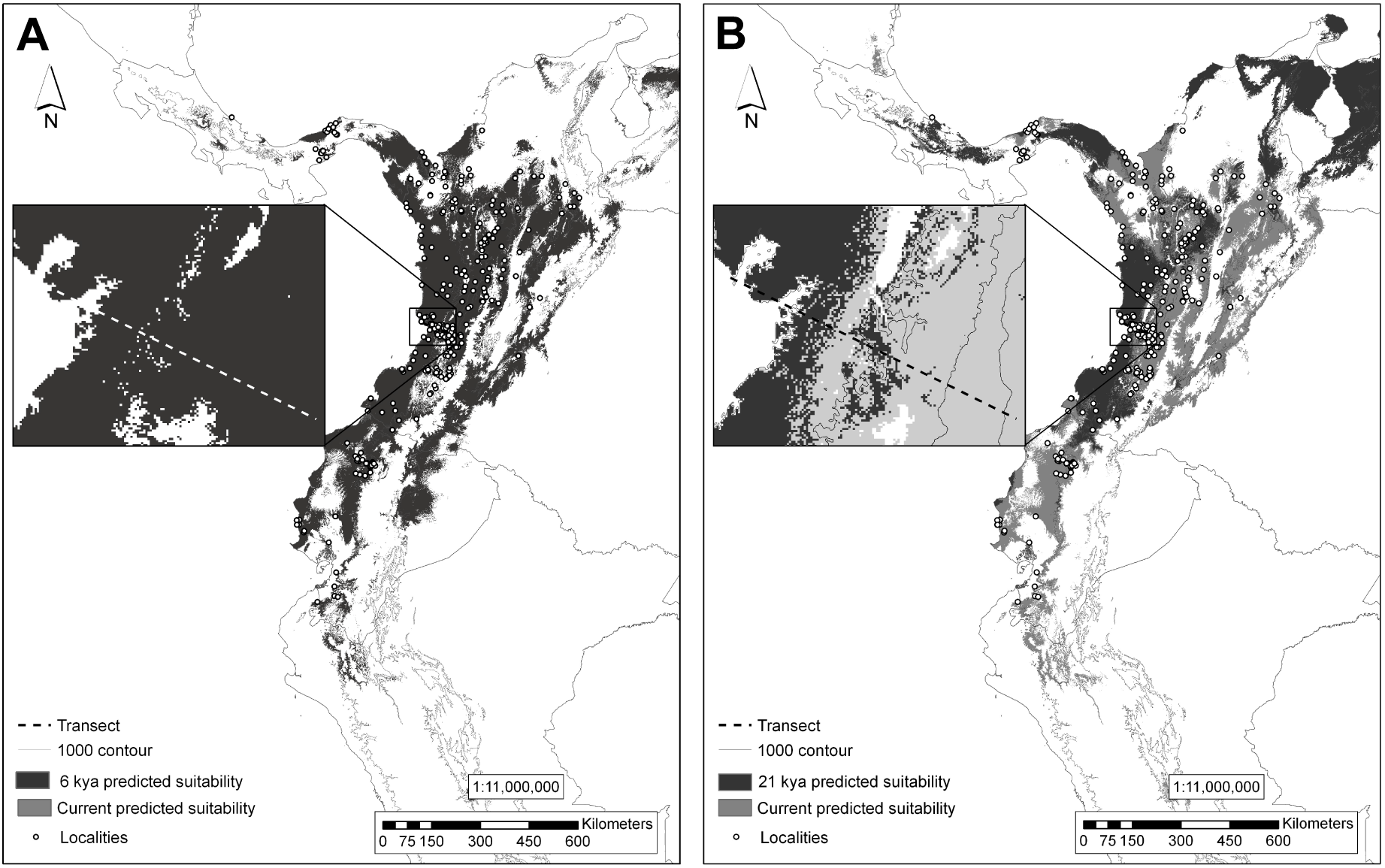
Potential distributions of *R. flammigerus* predicted by MaxEnt using climatic data. Dark gray areas show suitable climatic conditions for the occurrence of *flammigerus* and *icteronotus* (A) 6,000 years ago and (B) 21,000 years ago (LGM). Light gray depicts climatically suitable areas for their occurrence at present. Note the smaller predicted range during the LGM and that the two forms likely did not exhibit a continuous range along the transect (dotted line) at that time, relative to the more extensive and continuous range modeled for 6,000 y.a. and under current conditions. Dots represent localities used to construct the models.

### Genetic characterization

Overall, there was low genetic divergence between samples and genetic structure across the hybrid zone and among other localities was limited. Based on the recent samples for which we obtained long cytochrome *b* sequences, uncorrected mean sequence divergence within Colombia was only 0.3% (0-1.1%); samples from Ecuador and Colombia were 1.6% divergent, and samples from Panama and Colombia differed by only 0.4% on average. Except for a separation between samples from Ecuador and Colombia, relationships among haplotypes were not clearly resolved by the ML phylogenetic analysis, in which most nodes lacked bootstrap support and no clades associated with specific geographic regions or with plumage coloration were identified (Fig. 4). Among the long sequences (989 bp), there were 18 haplotypes with a total of 15 segregating sites in populations along our hybrid-zone transect. Among the 87 individuals from 1956 analyzed (210 bp), there were six haplotypes, with a total of nine segregating sites; uncorrected mean sequence divergence was only 0.4% (0-3.7%) and most (69) individuals shared a common haplotype. Relationships among haplotypes were not consistent with position along the hybrid zone (Fig. 4). For the same 210-bp region, there were seven haplotypes with six segregating sites in the 2010 specimens; clear structure with respect to position along the transect was not observed in the haplotype network (Fig. 4).

**Fig. 4.**
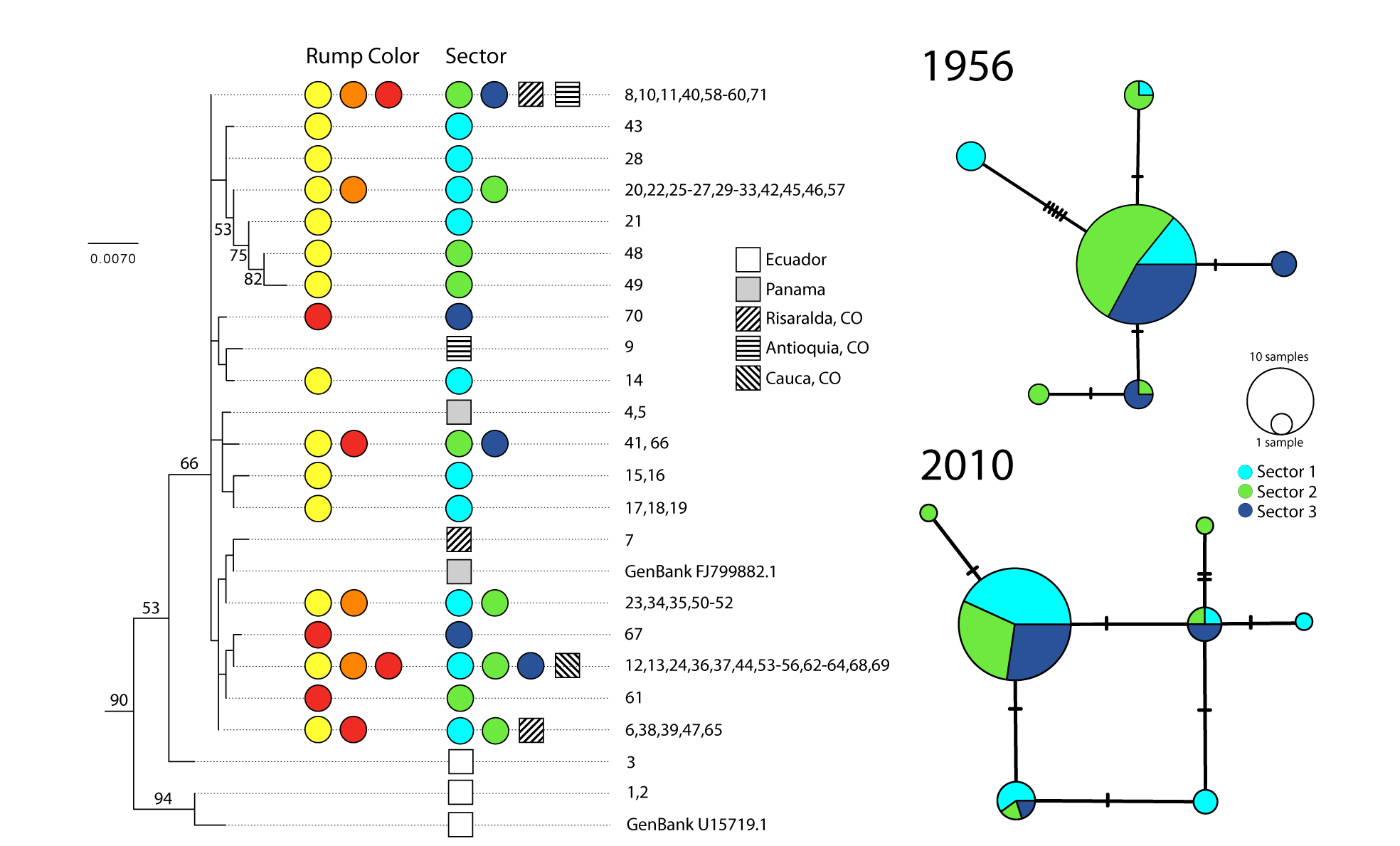
Genealogical relationships of specimens of *R. flammigerus* showing limited geographic structuring and relatively low levels of sequence divergence among haplotypes. The phylogenetic tree on the left depicts relationships among nearly complete sequences of the cytochrome *b* gene obtained for individuals from the hybrid zone and other localitites inferred using maximum-likelihood (outgroups not shown); bootstrap values on nodes are shown when ≥ 50%. Colored circles indicate a qualitative assessment of the rump color (yellow, orange and red as in individuals 1-6, 7-10 and 11-14 in Fig. 1, respectively) and location in the hybrid zone (cyan, sector 1; green, sector 2; dark blue, sector 3) of individuals from the study transect exhibiting each haplotype. The numbers correspond to specimen identifications in Table S1; all numbers refer to specimens from the hybrid-zone transect unless otherwise noted. Localities outside the transect in different provinces of Colombia (CO), Ecuador, and Panama are indicated with squares. Haplotype networks on the right focused on specimens from the hybrid zone show that genetic variation in 210 bp of the cytochrome *b* gene was not clearly consistent with position of individuals along the hybrid zone in 1956 (top) or in the present (bottom), although analyses of molecular variance suggest differences in patterns of genetic structure across time periods (see text); circle sizes are proportional to the number of individuals sharing each haplotype.

AMOVAs suggested that patterns of genetic structure across our study transect differ between specimens from 1956 and 2010, with considerably greater genetic structure among sectors in the 1956 data (Table 1). For the historical data, F_CT_ values were marginally significant (F_CT_ = 0.198, P = 0.011), indicating that a significant fraction of genetic variation (19%) was apportioned among sectors of the study transect. This was not the case for the present-day data, in which no genetic structure across the transect was detected (F_CT_ = 0.213, P = 0.111).

**Table 1.**
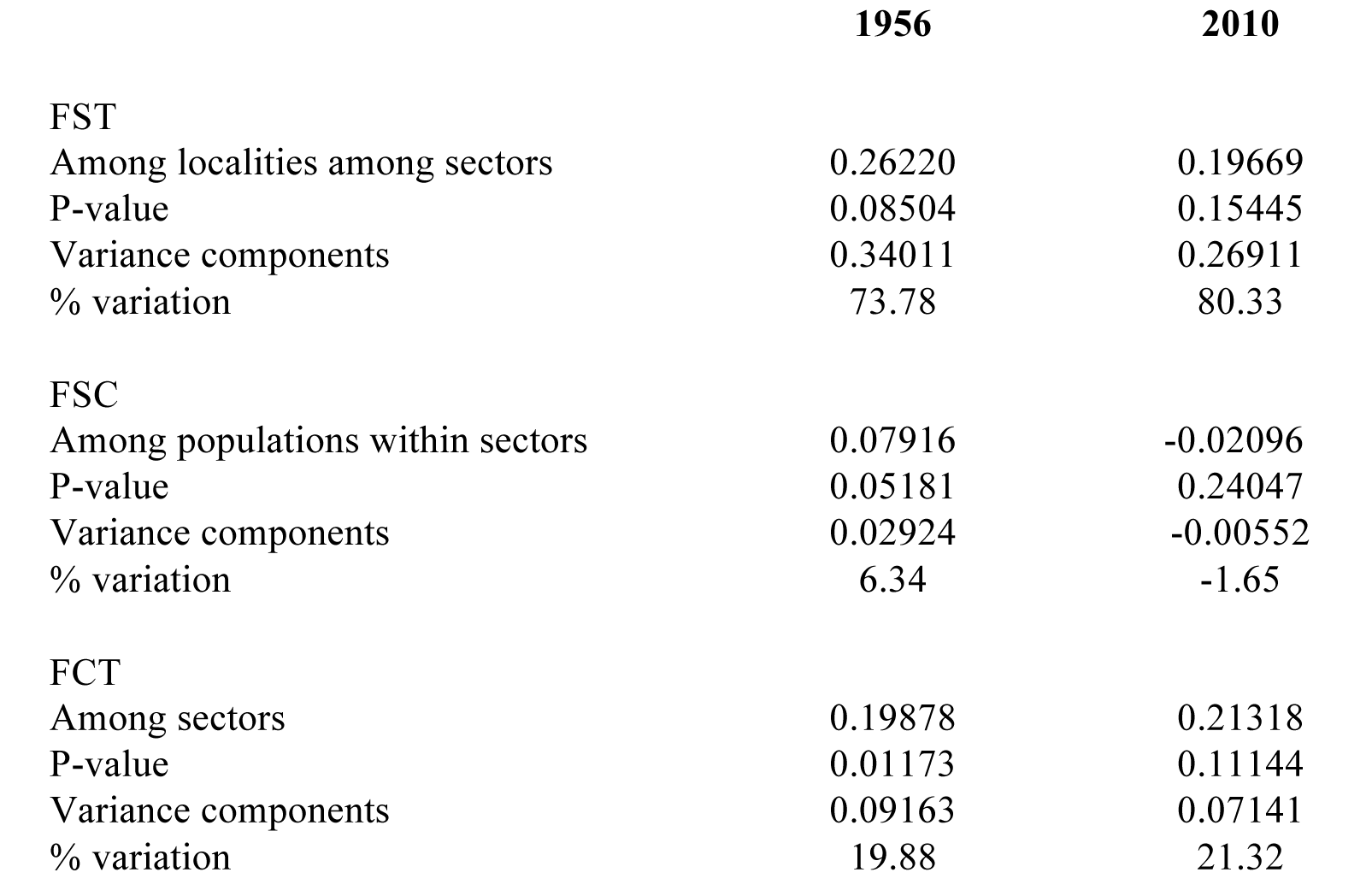
Population genetic structure in historical and recent specimens. Results are shown for analyses of molecular variance (AMOVA) based on DNA sequences of the cytochrome *b* mitochondrial gene for 87 individuals collected in 1956 and 58 individuals collected in 2010.

Although credibility intervals for population size in the Bayesian skyline plot were wide, this analysis suggested that populations show a genetic signature of demographic expansion (Fig. 5). Constant population size cannot be firmly rejected due to broad credibility intervals, but because the median value of the parameter “demographic.populationSizeChanges” differed from zero (median value = 1, 95% highest posterior density 0-2), the evidence points in the direction of population expansion rather than constant population size. This result is consistent with the hypothesis that the hybrid zone originated as a result of secondary contact following expansion of populations from formerly disjunct areas.

**Fig. 5.**
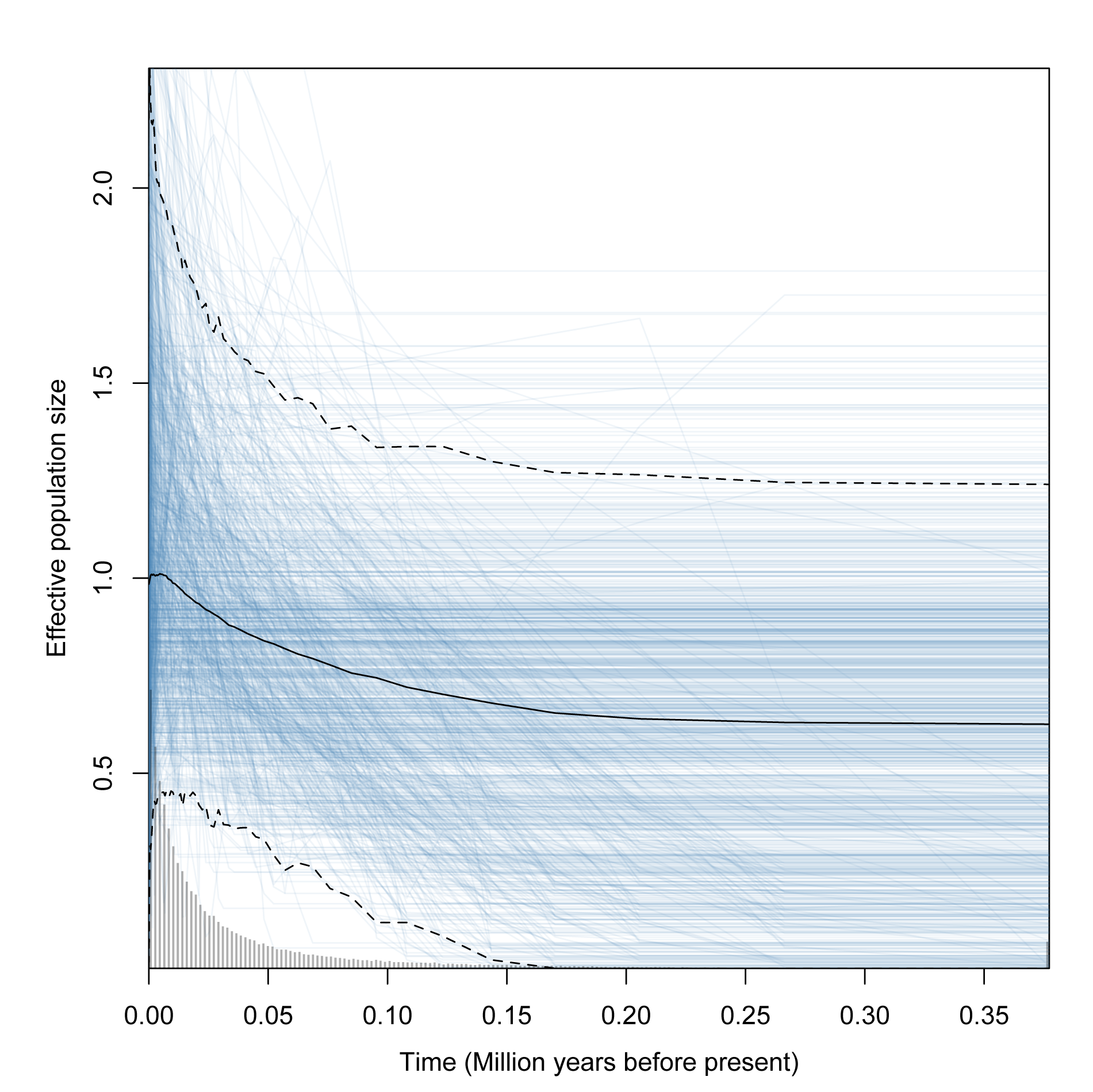
Estimates of population sizes over the last 400,000 years obtained using the extended Bayesian skyline plot method applied to cytochrome *b* sequence data suggest demographic expansion towards the present in *R. flammigerus*. Median and credibility interval values are shown in black solid line and dashed lines, respectively. Blue lines correspond to 1000 genealogies used to estimate the 95% highest posterior density of population sizes. Bars in the histogram are proportional to the number of genealogies with values in the specific time interval.

### Phenotypic characterization

Reduction of morphometric variation using PCA resulted in a first component (PC1) describing body size in both males and females. In both sexes, variables loading most heavily on this axis (which accounted for 24.3% of the variation in males and 34.7% in females) were tail length and wing chord. Thus, in the following we use PC1 as a general measurement of body size. We did not consider other principal component axes (e.g., PC2, on which bill dimensions loaded heavily) in additional analyses because they did not vary gradually across the hybrid zone.

Our estimates of cline parameters estimated using Hill functions were highly concordant with published estimates obtained using HZAR and Analyse in *Manacus* [65], especially for cline center (Table 2). Our estimates of cline width tended to differ more from published estimates, but in all cases the values estimated from the Hill function were within confidence intervals estimated by the other two algorithms. These results validate the use of the Hill function method to estimate cline parameters as described below.

**Table 2.**
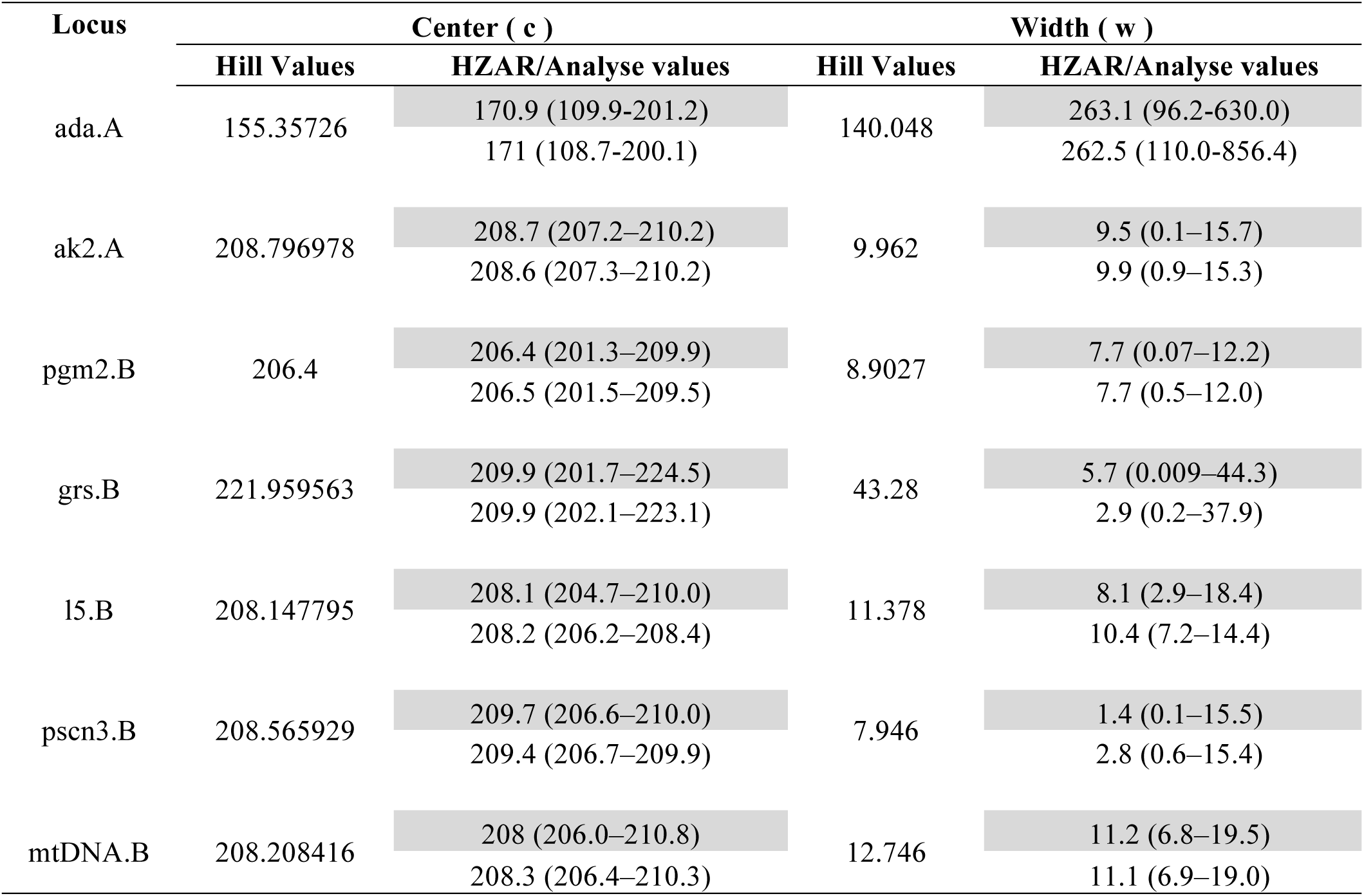
Validation of Hill-function methods as tools to estimate cline parameters. For each of seven genetic loci sampled across a *Manacus* hybrid zone in Panama, values of cline center and width estimated using Hill functions are similar to point estimates of these parameters obtained by a previous study [65] using cline-fitting algorithms HZAR (grey cells) and Analyse (white cells).

Morphological data for historical and recently collected male specimens provide evidence of clinal variation in body size (i.e. PC1) along the hybrid zone, with birds from localities to the west (*icteronotus*-type) being smaller than those from the east (*flammigerus*-type; Fig. 6). The Hill function method estimated that the center of the morphometric cline is currently located at ca. 76.7 km (mean across bootstrap replicates) from the coast extreme. The corresponding centers of the clines estimated using the pre-1911 and 1956-1986 specimens were at ca. 67.7 and 73.8 km, respectively, suggesting the zone has moved ca. 9 km over the past century (Table 3, Fig. 6, Fig.7, Fig. S1). Although differences among periods were significant (Tukey's post-hoc test; Table 4), bootstrap estimates of uncertainty around point estimates of cline centers overlapped broadly (Fig. 6G, Fig. 7).

**Fig. 6.**
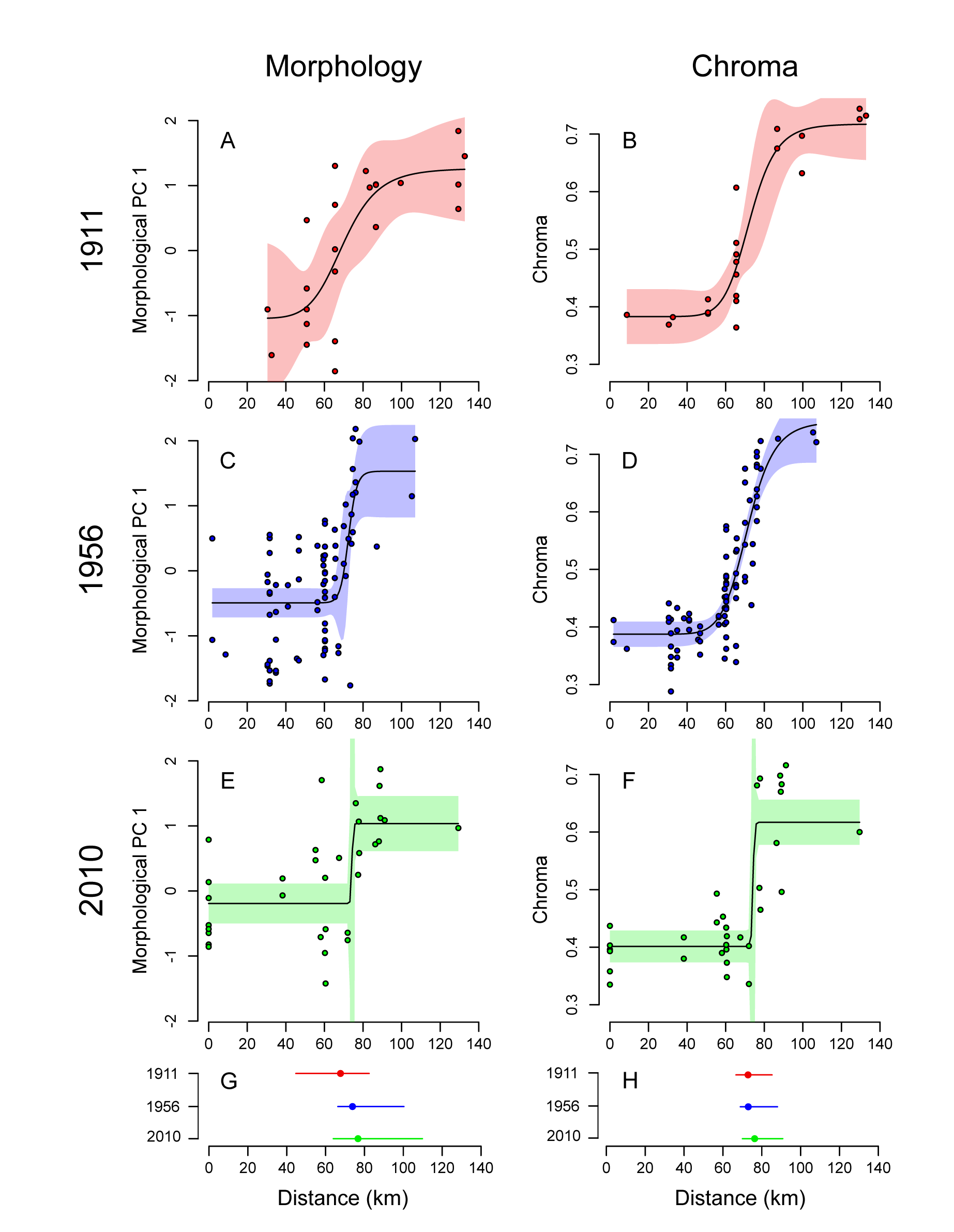
Variation in morphology and coloration of males over c. 130 km across the *Ramphocelus* hybrid zone in historical and recent specimens. **(A-F)** Circles are values for individual specimens; dark lines are clines for traits and periods in which parameters were estimated using Hill functions, with shading indicating 95% confidence intervals around cline estimates. **(G, H)** Estimates of cline centers at each time period and their confidence limits estimated using bootstrap samples, revealing coincident eastward movement of the hybrid zone over time as indicated by both traits (colors as in A-F).

**Figure 7.**
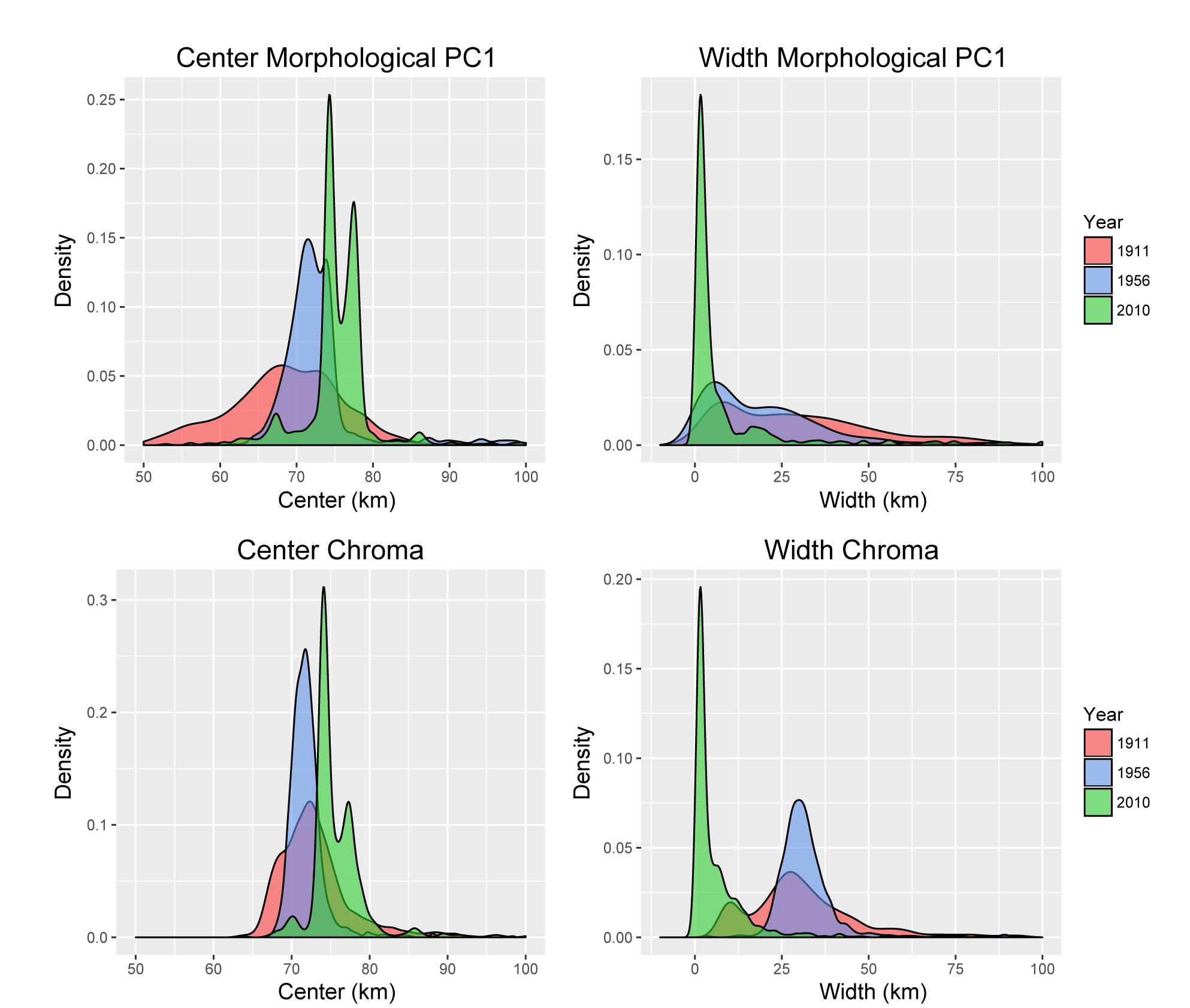
Probability density distibutions of cline center and cline width parameters estimated across 1,000 bootstrap samples of specimens for the 1911 (red), 1956 (blue), and 2010 (green)time periods. Both morphological PC1 and chroma show a shift in cline center and a reduction of cline width over time.

**Table 3.** Cline parameters estimated for morphological and coloration data in historical and recent specimens. The table shows mean values of cline centers and widths obtained from bootstrap samples of morphological variation (PC1) and plumage chroma for each time period based on data for male specimens. Values in parentheses correspond to the 95% confidence intervals. All values are given in kilometers, with cline centers measured as the distance along the transect from the Pacific coast extreme (Fig. 2). No confidence limit is given for the width parameter in morphology in 2010 because bootstrapping produced unrealistic parameters (see text).

|  |  | **Center ( c )** | **Width ( w )** |
| --- | --- | --- | --- |
| 1911 | Chroma | 72.92<br>(66.74-85.27) | 31.48<br>(8.78-78.69) |
|  | PC 1 | 67.67<br>(44.69-82.44) | 41.75<br>(3.61-165.4) |
| 1956 | Chroma | 73.04<br>(68.96-88.07) | 32.37<br>(21.14-60.48) |
|  | PC 1 | 73.85<br>(66.32-100.3) | 21.97<br>(0.84-86.43) |
| 2010 | Chroma | 76.33<br>(70.02-90.8) | 6.71<br>0.75-34.07 |
|  | PC 1 | 76.69<br>(63.95-109.84) | 33.12<br>- |

**Table 4.**
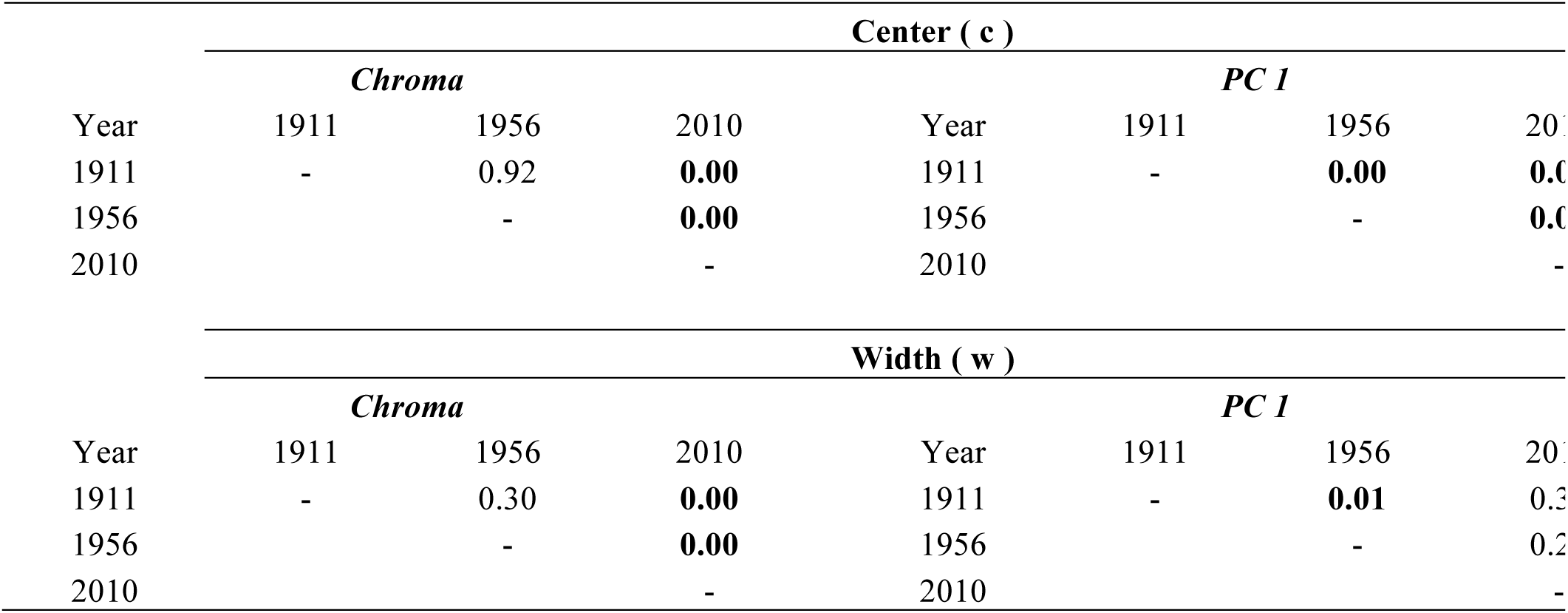
Results of post-hoc ANOVA comparing of clines center (c) and widths (w) among three time periods for morphological (PC1) and plumage coloration (chroma) data using the bootstrap distributions of parameters infered from Hill functions. Numbers correspond to *P* values of Tukey’s test, with those in bold indicating significant differences between periods.

Patterns of variation in color in space and time were similar to those observed for morphology. Of the three measurements of plumage coloration, chroma showed the clearest clinal pattern of variation, ranging from the yellow *icteronotus* to the redder *flammigerus* (Fig. 6). The Hill-function estimates of cline centers did not differ significantly between the pre-1911 (72.9 km) and 1956-1986 (73.0 km) periods, but the cline-center estimate for 2010 (76.3 km) was significantly different, suggesting a displacement of around 3 km in eastward direction (Table 3, Table 4, Fig. 6H, Fig. 7, Fig. S1).

Estimates of cline width were substantially more uncertain, but also appear to differ between the present and past. Cline width has declined across time, being wider in the pre-1911 (width 41.7 km) than in the 1956-1986 (22.0 km) and 2010 (33.1 km) specimens (Table 3, Fig. 7). Likewise, the estimated cline width in chroma at present was c. 6.7 km, but was wider in the past: 31.5 km in the pre-1911 specimens and 32.4 km 1956-1986 specimens (Table 3, Fig. 7).

Because sample sizes for females were much lower than those of males and because variation among female specimens across the hybrid zone was not clearly clinal (e.g., Fig. S2) we did not estimate cline parameters for female morphology and plumage measurements. Likewise, because measurements of plumage hue and brightness for males and females did not show clear clinal trends (data not shown), we did not attempt to estimate cline parameters for these traits.

## Discussion

Based on patterns of genetic variation, fossil pollen data, and ecological niche modeling, several studies in the north temperate zone indicate that the origin of hybrid zones can be explained as a result of population expansions from isolated refugia during the Quaternary [12,13,70]. Although a similar hypothesis was proposed to account for the origin of contact zones in tropical rainforest organisms [71,72], research on the origin of hybrid zones in the Neotropical region has been relatively limited [73,74]. Our niche models indicate that potential distributions of *R. f icteronotus* and *R. f flammigerus* were likely disjunct at the LGM (21,000 ya), but were potentially in contact by 6,000 ya. This scenario is supported by the historical demography analysis based on mtDNA sequence data, which indicates that populations have likely experienced significant range expansions, a pattern that awaits confirmation with multilocus data (see below) that should allow for lower uncertainty in estimates of population genetic parameters. Despite the broad credibility intervals around estimates of population size through time, taken together with results of niche modelling, the pattern observed in the skyline plot is consistent with a scenario in which forms likely diverged while isolated in each flank of the Cordillera Occidental (*icteronotus* in the Pacific lowlands and *flammigerus* in the Cauca Valley) and then expanded their distributions, presumably tracking the influence of Pleistocene climate change on vegetation [75,76]. A recent study also suggested that historical climatic changes likely promoted changes in the geographic distributions of lowland Neotropical birds which are presently separated by the Andes [77]. Our data also suggest that the divergence between the hybridizing *Ramphocelus* populations likely occurred in the Pleistocene, as indicated by low levels of mtDNA divergence suggesting recent differentiation. However, because Quaternary climatic oscillations started well before the LGM [78], it is possible that distribution ranges became disjunct and reconnected repeatedly at various times throughout the Pleistocene.

In contrast to our proposed scenario suggesting the origin of the *Ramphocelus* hybrid zone may date to at least 6,000 before present, Sibley [25] hypothesized that contact between *flammigerus* and *icteronotus* resulted from recent anthropogenic deforestation and expansion of crops creating scrub and second-growth habitats, which are favored by these tanagers over dense rain forest. Although our analyses suggest that climatic conditions were suitable for contact between these forms thousands of years prior to human alterations in the area, it is likely that anthropogenic activities have facilitated contact between them, possibly leading to an increased incidence of hybridization in recent times.

A recent study on a hybrid zone between *Heliconius* butterflies located in the same geographic region where we studied hybridization in *Ramphocelus* also provided evidence consistent with the hypothesis of origin via secondary contact [79]. Because there are additional documented cases of hybridization in the same general area of southwestern Colombia (e.g., other *Heliconius* butterfiles [80], *Oophaga* poison frogs [81]), work on the history of the region is necessary to better understand the origin and maintenance of hybrid zones across taxa [12,15].

Our analyses are consistent with Sibley's [25] overall characterization of the *Ramphocelus* hybrid zone: there is clinal variation in coloration and body size, with males exhibiting clearer trends than females (see also [29]). With the caveat that uncertainty around parameter estimates is broad, two main additional insights are provided by our cline analyses. First, our data consistently indicate that for each period, clines for morphology and chroma are coincident (i.e., they have equal or very similar centers). Second, variation in morphological PC1 and chroma followed the same trend over time: for both traits, clines moved to the east a few kilometers and became narrower from the past to the present.

The coincidence of cline centers for different characters and their apparent concordance in width in each of the three periods is consistent with a tension-zone model. This is further supported by the width estimated for character clines. Assuming that the hybrid zone originated at least 6,000 ya according to climate-based models, that generation time in *Ramphocelus* is 1-2 years, and that dispersal distances per generation lie somewhere between 1 and 20 km, the expected width of clines under a neutral diffusion model (equations in [7,82]) would be between c. 25 km and more than 500 km. This is much wider than what we estimated (Table 2), which suggests some form of selection is acting. Alternatively, narrow clines may be a result of a more recent origin of the hybrid zone than implied by climate data. However, even if one assumes that the hybrid zone is as young as proposed by Sibley [25], the estimated clines appear narrower than expected under neutral diffusion (c. 4-75 km). In sum, our results suggest that the *Ramphocelus* hybrid zone is likely a tension zone, maintained by a balance between dispersal and a genome-wide barrier to gene flow [1]. This interpretation contrasts with Sibley's [25] conclusions that “gene exchange appears to be unimpeded” and that there is “no evidence of selection against the hybrids”. Future studies should further evaluate our tension-zone hypothesis by examining survival and mating success of intermediate phenotypes resulting from hybridization relative to parental types.

That cline centers do not appear to be coincident across different periods for each of the characters suggests movement of the *Ramphocelus* hybrid zone, but this interpretation needs to be tempered because samples were not taken at exactly the same localities at each time period and because the centers we estimated in some cases had wide support limits. However, our field observations indicate that the current patterns of variation are, in fact, different from those described by Sibley [25]. Specifically, we frequently observed yellow-rumped individuals at Salado and Queremal, where Sibley did not report any, and we also have occasional records of yellow-rumped individuals near Cali, the eastern extreme of the transect. Thus, our quantitative analyses and field observations suggest that the *icteronotus* phenotype (yellow rump and smaller body size) has indeed extended to the east. Several previous studies have also reported on moving tension zones [83-85] although others documented spatial stability [86,87].

A possible explanation for the concurrent movement of clines for different characters in hybrid zones is competitive advantage of one phenotype mediated by aggression or by sexual selection [18]. Thus, movement of the *Ramphocelus* hybrid zone may have been driven by competition or sexual selection favoring the *icteronotus* phenotype. We believe it is unlikely that aggressive superiority of *icteronotus* explains this pattern as shown in other studies of hybrid zones owing to its smaller body size, although we note that in a *Manacus* (Pipridae) hybrid zone in Panama, the smaller *M. vitellinus* is more aggressive than the larger *M. candei* [88]. Thus, it would be of interest to study mating patterns in the field and to conduct mate-choice experiments to determine whether the *icteronotus* phenotype has increased reproductive success as a result of sexual selection via male-male dominance or female choice [89,90].

If sexual selection is based on carotenoid plumage color, which presumably is strongly influenced by the environment [32], then it is somewhat puzzling that coloration seems to have moved across the hybrid zone in concert with morphometric variation, which presumably has high heritability and is not involved in mate choice. It is possible, however, that the ability to obtain, accumulate and metabolize carotenoids has a heritable genetic basis [91] and that the fitness advantages it confers may be linked to genes involved in reproductive isolation [92]. If this were the case in *R. flammigerus*, then it would represent a plausible explanation for the movement of coloration in concert with others traits.

An alternative explanation for zone movement was proposed by Sibley [25], who predicted that the *icteronotus* phenotype would introgress across the hybrid zone as a consequence of increased gene flow from coastal to interior populations resulting from larger populations sizes in the former. This could be studied in the future with multilocus estimates of effective population sizes and of the magnitude of gene flow in both directions [93]. In addition, if deforestation has indeed resulted in increases in population size as hypothesized by Sibley [25], then movement of the hybrid zone may also partly reflect anthropogenic influences. Because hybrid zones tend to become entrapped in areas of low population densities acting as sinks for migration [1,87], any changes in population size related to habitat modification may have partly facilitated the observed movement of the *Ramphocelus* hybrid zone.

We note that the inferred movement of the hybrid zone involves not only a west-east displacement, but also a shift in its center to higher elevations in the Andes. Because the elevational ranges of tropical birds may shift upslope in response to global warming [94], it is also possible that the movement of the hybrid zone is related to climatic change over the past few decades [18,22,95,96], as recently documented for hybrid zones between woodpecker (Picidae) and chickadee (Paridae) species in North America [97,98].

In contrast to multiple studies on hybridization in birds finding significant mtDNA divergence between populations located away from the center of hybrid zones and clinal variation in haplotype frequencies across them (e.g., [40,44,54,73,86,99-103]), mtDNA variation was not geographically structured in our study system, a likely consequence of recent divergence of the hybridizing populations or of high levels of introgression. Because no clinal variation was observed across the *Ramphocelus* hybrid zone, we were unable estimate cline parameters for different time periods to examine zone movement as done in a few studies on hybrid zones examining genetic variation in specimens collected at different times [23,85,101,104]. However, our analyses revealing more limited genetic structure across the zone as indicated by F-statistics in 2010 relative to 1956 provide some evidence that patterns of genetic variation may have not been stable over time. We hypothesize that these results may reflect an increase in introgression of mtDNA across our study transect since Sibley's [25] time, but we acknowledge that because we only assayed a small fragment of mtDNA and because sampling was not spatially even between time periods, drawing any conclusion at this time would be premature. Nonetheless, our preliminary genetic data suggest that assessing temporal variation in spatial genetic structure using genome-wide markers (cf. [43]) would represent a fruitful avenue to better understand the ecological and evolutionary forces at work in this moving hybrid zone. Also, if shallow mtDNA divergence reflects overall patterns in neutral divergence, then the *Ramphocelus* hybrid zone appears especially well-suited for analyses of variation in functionally important genes, which may provide important insights about the genetic basis of phenotypic variation [105,106].

Finally, our study serves to illustrate that Hill functions and bootstrap resampling are useful statistical tools to characterize patterns of variation across hybrid zones. In contrast to other studies [e.g., 23], our work was not systematically designed to examine changes in the structure of the *Ramphocelus* hybrid zone over time, with repeated sampling of multiple specimens at fixed locations; rather, specimens were collected by numerous researchers, often opportunistically, for different purposes, and with no common sampling schemes at different times. Therefore, we did not feel confident in assigning individuals to discrete localities as required for cline-fitting algorithms which rely on calculating means and variances for samples of specimens from the same sites [65-67]. Our reanalysis of a previously published data set suggests that cline parameters estimated using Hill functions mirror closely those estimated by software commonly employed in studies of hybrid zones, suggesting such functions are useful alternatives when methods designed to study clines cannot be employed due to the nature of the available data. Although Hill functions have been used mainly to model dose-response curves, in principle they can be used to describe any relationship between variables that follows a sigmoid shape. In addition to not requiring data to be grouped in localities (i.e. each specimen conserves its position along sampling transects), the approach we followed does not assume normality in the data and can be run with low computational requeriments and relatively small sample sizes. On the other hand, we suspected that due to variation in the spatial distribution and number of specimens over time, inferences about temporal changes in the structure of the hybrid zone could be compromised by unequal sampling. However, by analyzing variation in bootstrap samples of specimens, we were able to account for sampling effects, finding that despite inconsistent sampling over time, there is sufficient signal in our data to document temporal changes in spatial patterns of phenotypic variation. Similar approaches could be employed to analyse temporal changes in spatial variation in genetic or phenotypic traits in other systems in which museum specimens have not been systematically collected over time and space but nonetheless contain rich historical information.

## Conclusions

Models of potential historical distributions based on climatic data and genetic signatures of demographic expansion suggested that the *Ramphocelus* hybrid zone we studied originated following secondary contact between populations that expanded their ranges out of isolated areas in the Quaternary, likely as a consequence of climatic change. Concordant patterns of variation in phenotypic characters across the hybrid zone and its narrow extent are suggestive of a tension zone, maintained by a balance between dispersal and selection against hybrids. In addition, our estimates of phenotypic cline parameters obtained using specimens collected over nearly a century revealed that the zone has moved to the east and to higher elevations, and has likely become narrower. These observed changes in the hybrid zone may be a result of sexual selection, asymmetric gene flow, or environmental change. Our data represent a baseline for a variety of ecological, behavioral, and genetic studies that could shed further light on the forces involved in speciation and hybrid-zone dynamics in this system.

## Authors’ Contributions

AMR and CDC conceived the study. AMR collected and measured specimens, conducted molecular genetic work, and performed initial analyses. MDC conducted molecular work on historical specimens. AMR and EAT analyzed data with supervision from CDC. AMR and CDC wrote the manuscript, with input from EAT and MDC. All authors read, commented on, and approved the final manuscript.

## Acknowledgments

The Facultad de Ciencias at Universidad de los Andes, an American Ornithologists’ Union Research Award, and a Collection Study Grant from the Department of Ornithology at the American Museum of Natural History provided funding for this study. The Ministerio de Ambiente, Vivienda y Desarrollo Territorial of Colombia provided permits for research, collecting, and accessing genetics resources (RGE0014). F. Ayerbe, J. P. Lopéz, J. P. Goméz, N. Gutiérrez, S. Gonzalez, P. Tovar, F. Gonzalez, C. Wagner, J. Sandoval, the Luna Family, and the Castaño family helped in the field. For assistance with analyses, we are grateful to J. M. Cordovez, C. Ritz, C. Palacios, L. N. Céspedes, E. Derryberry, N. Gutiérrez, J. L. Parra, C. Pedraza, and E. Valderrama. We thank the Instituto de Génetica at Universidad de Los Andes (M. Linares) for access to facilities, and the curators and collection managers at institutions who allowed us to examine specimens or provided information or tissue samples for genetic analyses: American Museum of Natural History (J. Cracraft, P. Sweet), Cornell University Museum of Vertebrates (K. Botswick, I. Lovette), Instituto de Ciencias Naturales at Universidad Nacional de Colombia (F. G. Stiles), Louisiana State University Museum of Natural Science (R. Brumfield), Universidad del Cauca (M. D. P. Rivas), Universidad del Pacífico (A. Quintero), and Universidad del Valle (H. Alvarez-López).

---

**Figure S1.**
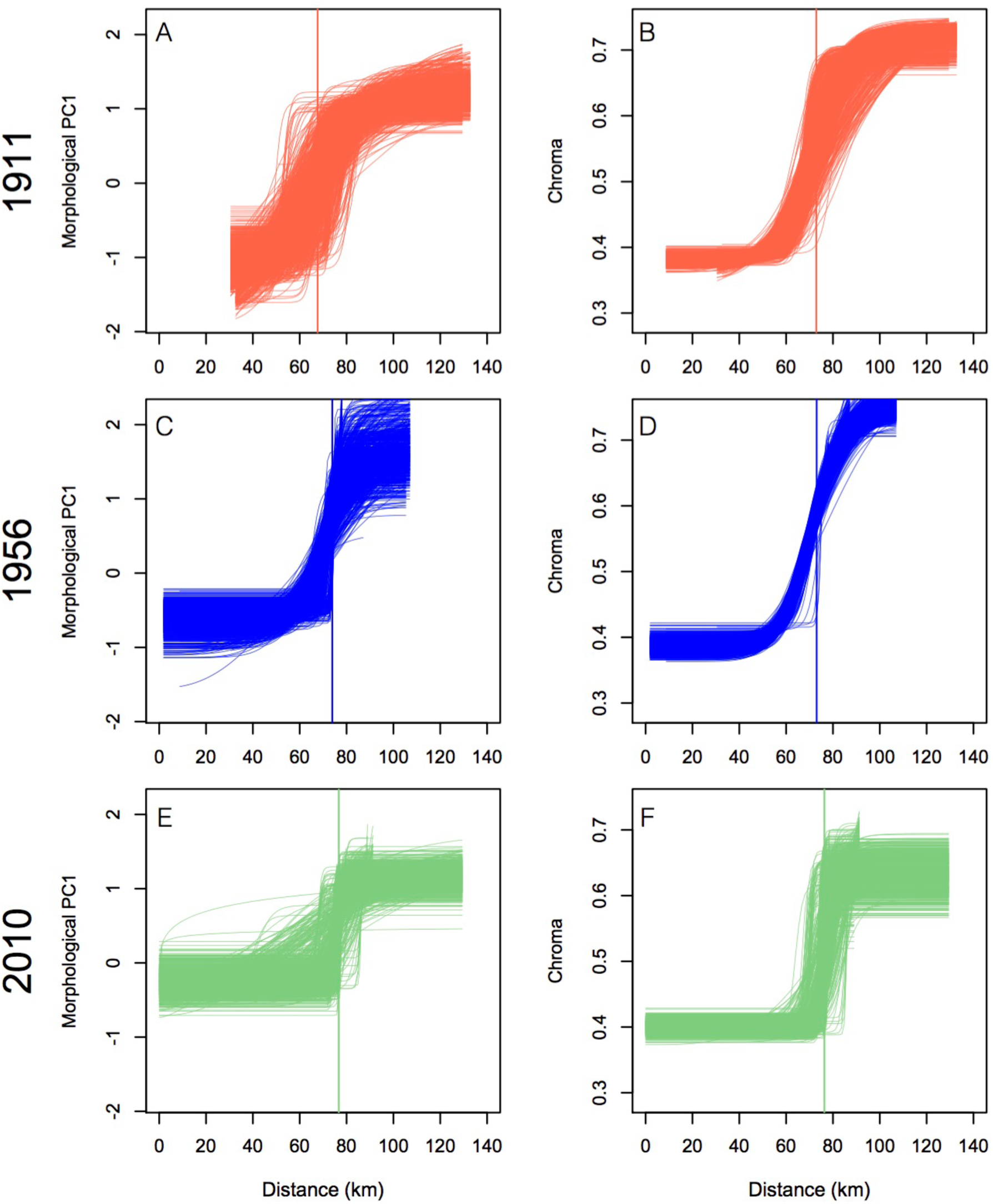
Clines for morphological and plumage data estimated across 1,000 bootstrap samples. The vertical line in each plot corresponds to the mean value of cline centers for the 1911 (A,B), 1956 (C,D) and 2010 (E,F) periods.

**Figure S2.**
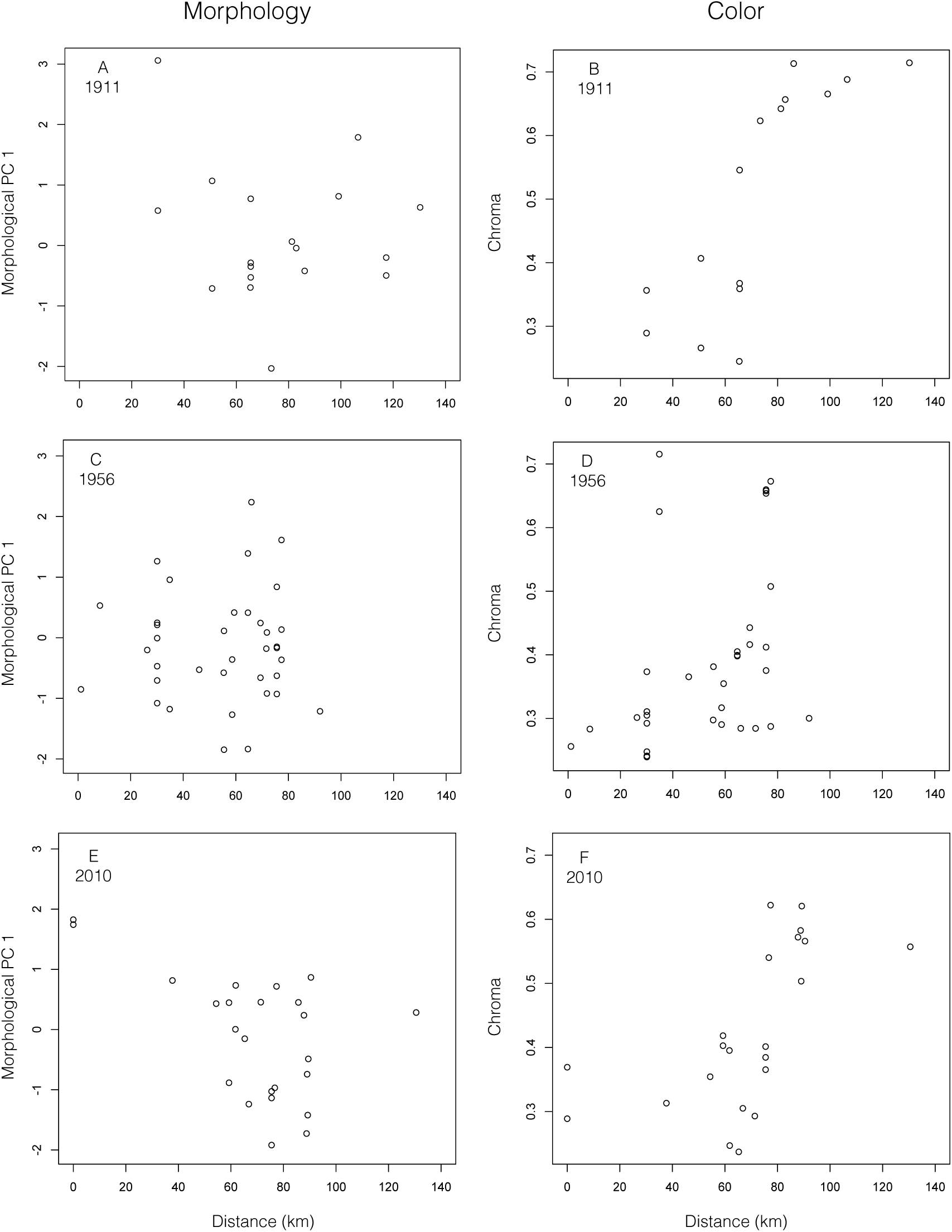
Variation in morphology (morphological PC1) and plumage chroma in female specimens across the *Ramphocelus flammigerus* hybrid zone in southwestern Colombia. Data are shown separately for historical specimens (A,B: 1911; C,D: 1956) and recent specimens (E,F: 2010).

**Table S1.** Information on the samples of *Ramphocelus flammigerus* included in the study, including identification number (ID), locality, museum catalogue number, geographical coordinates, and GenBank accession numbers for cytb sequences. Museum acronyms are as follows: LSUMZ: Louisiana State University Museum of Natural Science; ANDES-O: Museo de Historia Natural, Universidad de los Andes CUMV: Cornell University Museum of Vertebrates, Cornell University.

| **Id** | **Locality** | **Catalogue No.** | **Latitude** | **Longitude** | **Accession No.** |
| --- | --- | --- | --- | --- | --- |
| 1 | Ecuador, Esmeraldas Province El Placer | LSUMZ B-12014 | 0.867 | -78.550 | KR869689 |
| 2 | Ecuador, Esmeraldas Province, El Placer | LSUMZ B-12017 | 0.867 | -78.550 | KR869688 |
| 3 | Ecuador, Pichincha Province 5 km | LSUMZ B-35022 | 0.017 | -78.883 | KR869690 |
| 4 | Panamá, Colon Province: Achioté Road | LSUMZ B-28758 | 9.223 | -80.019 | KR869691 |
| 5 | Panamá, Darien Province | LSUMZ B-52947 | 8.083 | -77.883 | KR869692 |
| 6 | Colombia, Risaralda, Pueblo Rico | ANDES-O AMR106 | 5.231 | -76.084 | KR869693 |
| 7 | Colombia, Risaralda, Pueblo Rico | ANDES-O AMR107 | 5.231 | -76.084 | KR869694 |
| 8 | Colombia, Risaralda, Pueblo Rico | ANDES-O NGP93 | 5.235 | -76.085 | KR869695 |
| 9 | Colombia, Antioquia, San Roque | ANDES-O AMR118 | 6.384 | -74.992 | KR869696 |
| 10 | Colombia, Antioquia, Cocorná | ANDES-O AMR55 | 6.027 | -75.144 | KR869697 |
| 11 | Colombia, Antioquia, Cocorná | ANDES-O AMR56 | 6.027 | -75.144 | KR869698 |
| 12 | Colombia, Cauca, Popayán | ANDES-O AMR83 | 2.477 | -76.575 | KR869700 |
| 13 | Colombia, Cauca, Popayán | ANDES-O AMR84 | 2.477 | -76.575 | KR869699 |
| 14 | Colombia, Valle del Cauca, Buenaventura, La Barra | ANDES-O NGP101 | 3.963 | -77.379 | KR817417 |
| 15 | Colombia, Valle del Cauca, Buenaventura, La Barra | ANDES-O NGP99 | 3.963 | -77.379 | KR817418 |
| 16 | Colombia, Valle del Cauca, Buenaventura, La Barra | ANDES-O AFR29 | 3.963 | -77.379 | KR817419 |
| 17 | Colombia, Valle del Cauca, Ladrilleros, Vía la Barra | ANDES-O AMR39 | 3.946 | -77.366 | KR817423 |
| 18 | Colombia, Valle del Cauca, Buenaventura, La Barra | ANDES-O AMR86 | 3.963 | -77.379 | KR817420 |
| 19 | Colombia, Valle del Cauca, Buenaventura, Ladrilleros | ANDES-O AMR40 | 3.944 | -77.366 | KR817422 |
| 20 | Colombia, Valle del Cauca, B/tura, La Barra | ANDES-O AMR112 | 3.963 | -77.379 | KR817459 |
| 21 | Colombia, Valle del Cauca, Buenaventura, La Barra | ANDES-O AMR111 | 3.959 | -77.376 | KR817428 |
| 22 | Colombia, Valle del Cauca, Buenaventura, La Barra | ANDES-O AMR87 | 3.964 | -77.379 | KR817473 |
| 23 | Colombia, Valle del Cauca, Buenaventura, Ladrilleros | ANDES-O AMR41 | 3.946 | -77.366 | KR817440 |
| 24 | Colombia, Valle del Cauca, Buenaventura | ANDES-O AMR105 | 3.848 | -77.000 | KR817442 |
| 25 | Colombia, Valle del Cauca, Buenaventura | ANDES-O AMR103 | 3.848 | -77.000 | KR817460 |
| 26 | Colombia, Valle del Cauca, Buenaventura | ANDES-O AMR104 | 3.848 | -77.000 | KR817464 |
| 27 | Colombia, Valle del Cauca, Dagua, El Placer | ANDES-O AMR95 | 3.591 | -76.814 | KR817466 |
| 28 | Colombia, Valle del Cauca, Dagua, Alto Anchicayá | ANDES-O AMR47 | 3.575 | -76.855 | KR817462 |
| 29 | Colombia, Valle del Cauca, Dagua, Alto Anchicayá | ANDES-O AMR69 | 3.573 | -76.879 | KR817472 |
| 30 | Colombia, Valle del Cauca, Dagua, Bajo | ANDES-O NGP28 | 3.609 | -76.918 | KR817463 |
| 31 | Colombia, Valle del Cauca, Dagua, Alto Anchicayá | ANDES-O AMR68 | 3.535 | -76.871 | KR817471 |
| 32 | Colombia, Valle del Cauca, Dagua, Bajo Anchicayá | ANDES-O AMR64 | 3.609 | -76.918 | KR817470 |
| 33 | Colombia, Valle del Cauca, Dagua, Alto Anchicayá | ANDES-O AMR45 | 3.574 | -76.878 | KR817474 |
| 34 | Colombia, Valle del Cauca, Dagua, Alto Anchicayá | ANDES-O NGP27 | 3.576 | -76.880 | KR817435 |
| 35 | Colombia, Valle del Cauca, Dagua, Alto Anchicayá | ANDES-O AMR66 | 3.576 | -76.880 | KR817439 |
| 36 | Colombia, Valle del Cauca, Dagua, Alto Anchicayá | ANDES-O NGP30 | 3.576 | -76.880 | KR817445 |
| 37 | Colombia, Valle del Cauca, Dagua, Bajo Anchicayá | ANDES-O AMR65 | 3.609 | -76.918 | KR817446 |
| 38 | Colombia, Valle del Cauca, Buenaventura, Los Tubos | ANDES-O AMR42 | 3.845 | -76.793 | KR817433 |
| 39 | Colombia, Valle del Cauca, Buenaventura, La Delfina | ANDES-O AMR43 | 3.836 | -76.791 | KR817432 |
| 40 | Colombia, Valle del Cauca, Dagua, Queremal | ANDES-O AMR102 | 3.595 | -76.795 | KR817455 |
| 41 | Colombia, Valle del Cauca, Dagua, La Elsa | ANDES-O AMR98 | 3.576 | -76.756 | KR817424 |
| 42 | Colombia, Valle del Cauca, Dagua, Alto Anchicayá | ANDES-O NGP29 | 3.573 | -76.878 | KR817465 |
| 43 | Colombia, Valle del Cauca, Dagua, Alto Anchicayá | ANDES-O AMR63 | 3.576 | -76.879 | KR817430 |
| 44 | Colombia, Valle del Cauca, Dagua, La Elsa | ANDES-O AMR97 | 3.576 | -76.756 | KR817443 |
| 45 | Colombia, Valle del Cauca, Dagua, Alto Anchicayá | ANDES-O AMR48 | 3.575 | -76.879 | KR817468 |
| 46 | Colombia, Valle del Cauca, Dagua, Alto Anchicayá | ANDES-O AMR49 | 3.574 | -76.878 | KR817469 |
| 47 | Colombia, Valle del Cauca, Dagua, Alto Anchicayá | ANDES-O AMR46 | 3.575 | -76.879 | KR817431 |
| 48 | Colombia, Valle del Cauca, Dagua, Queremal | ANDES-O AMR70 | 3.528 | -76.719 | KR817421 |
| 49 | Colombia, Valle del Cauca, Dagua, Queremal | ANDES-O AMR74 | 3.527 | -76.713 | KR817426 |
| 50 | Colombia, Valle del Cauca, Dagua, Queremal | ANDES-O AMR78 | 3.526 | -76.733 | KR817436 |
| 51 | Colombia, Valle del Cauca, Dagua, Queremal | ANDES-O AMR75 | 3.526 | -76.733 | KR817437 |
| 52 | Colombia, Valle del Cauca, Dagua, Queremal | ANDES-O AMR66 | 3.576 | -76.880 | KR817439 |
| 53 | Colombia, Valle del Cauca, Dagua, Salado | ANDES-O AMR73 | 3.571 | -76.697 | KR817451 |
| 54 | Colombia, Valle del Cauca, Dagua, Queremal | ANDES-O AMR76 | 3.526 | -76.733 | KR817447 |
| 55 | Colombia, Valle del Cauca, Dagua, Queremal | ANDES-O AMR77 | 3.526 | -76.733 | KR817452 |
| 56 | Colombia, Valle del Cauca, Dagua, Queremal | ANDES-O AMR72 | 3.526 | -76.719 | KR817444 |
| 57 | Colombia, Valle del Cauca, Cali, La Leonera | ANDES-O AMR85 | 3.455 | -76.635 | KR817467 |
| 58 | Colombia, Valle del Cauca, La Cumbre | ANDES-O AMR37 | 3.575 | -76.572 | KR817457 |
| 59 | Colombia, Valle del Cauca, La Cumbre, Chicoral | ANDES-O AMR36 | 3.566 | -76.581 | KR817458 |
| 60 | Colombia, Valle del Cauca, Dagua, San Bernardo | ANDES-O AMR52 | 3.474 | -76.647 | KR817456 |
| 61 | Colombia, Valle del Cauca, La Cumbre, Chicoral | ANDES-O AMR24 | 3.566 | -76.581 | KR817461 |
| 62 | Colombia, Valle del Cauca, La Cumbre, Chicoral | ANDES-O AMR25 | 3.566 | -76.581 | KR817453 |
| 63 | Colombia, Valle del Cauca, Cali, Peñas Blancas | ANDES-O AMR51 | 3.428 | -76.643 | KR817449 |
| 64 | Colombia, Valle del Cauca, Cali, Peñas Blancas | ANDES-O AMR50 | 3.428 | -76.643 | KR817448 |
| 65 | Colombia, Valle del Cauca, La Cumbre, Chicoral | ANDES-O AMR38 | 3.571 | -76.575 | KR817434 |
| 66 | Colombia, Valle del Cauca, Cali, Andes | ANDES-O NGP102 | 3.422 | -76.617 | KR817425 |
| 67 | Colombia, Valle del Cauca, Cali, Andes | ANDES-O AMR114 | 3.424 | -76.617 | KR817429 |
| 68 | Colombia, Valle del Cauca, Cali, La Elvira | ANDES-O AMR53 | 3.526 | -76.594 | KR817450 |
| 69 | Colombia, Valle del Cauca, Cali, Andes | ANDES-O AMR113 | 3.422 | -76.617 | KR817441 |
| 70 | Colombia, Valle del Cauca, Palmira, La Buitrera | ANDES-O AMR115 | 3.476 | -76.192 | KR817427 |
| 71 | Colombia, Valle del Cauca, Palmira, La Buitrera | ANDES-O AN006 | 3.471 | -76.189 | KR817454 |
| 72 | Colombia, Valle del Cauca, Buenaventura | CUMV 27061 | 3.794 | -76.993 | KR817483 |
| 73 | Colombia, Valle del Cauca, Buenaventura | CUMV 27062 | 3.794 | -76.993 | KR817542 |
| 74 | Colombia, Valle del Cauca, Buenaventura | CUMV 27063 | 3.794 | -76.993 | KR817543 |
| 75 | Colombia, Valle del Cauca, Buenaventura | CUMV 27064 | 3.794 | -76.993 | KR817544 |
| 76 | Colombia, Valle del Cauca, Buenaventura | CUMV 27066 | 3.844 | -77.000 | KR817476 |
| 77 | Colombia, Valle del Cauca, Buenaventura | CUMV 27067 | 3.794 | -76.993 | KR817477 |
| 78 | Colombia, Valle del Cauca, Buenaventura, Río Anchicaya | CUMV 27037 | 3.614 | -76.905 | KR817528 |
| 79 | Colombia, Valle del Cauca, Buenaventura, Río Anchicaya | CUMV 27038 | 3.614 | -76.905 | KR817529 |
| 80 | Colombia, Valle del Cauca, Buenaventura, Río Anchicaya | CUMV 27039 | 3.614 | -76.905 | KR817475 |
| 81 | Colombia, Valle del Cauca, Buenaventura, Río Anchicaya | CUMV 27040 | 3.614 | -76.905 | KR817530 |
| 82 | Colombia, Valle del Cauca, Buenaventura, Río Anchicaya | CUMV 27041 | 3.614 | -76.905 | KR817531 |
| 83 | Colombia, Valle del Cauca, Buenaventura, Río Anchicaya | CUMV 27042 | 3.614 | -76.905 | KR817491 |
| 84 | Colombia, Valle del Cauca, Buenaventura, Zabaletas | CUMV 27069 | 3.745 | -76.953 | KR817478 |
| 85 | Colombia, Valle del Cauca, Buenaventura, Zabaletas | CUMV 27072 | 3.745 | -76.953 | KR817546 |
| 86 | Colombia, Valle del Cauca, Buenaventura, Zabaletas | CUMV 27073 | 3.745 | -76.953 | KR817547 |
| 87 | Colombia, Valle del Cauca, Buenaventura, El Placer | CUMV 27013 | 3.606 | -76.868 | KR817513 |
| 88 | Colombia, Valle del Cauca, Buenaventura, El Placer | CUMV 27015 | 3.606 | -76.868 | KR817514 |
| 89 | Colombia, Valle del Cauca, Buenaventura, El Placer | CUMV 27016 | 3.606 | -76.868 | KR817515 |
| 90 | Colombia, Valle del Cauca, Buenaventura, El Placer | CUMV 27017 | 3.606 | -76.868 | KR817479 |
| 91 | Colombia, Valle del Cauca, Buenaventura, El Placer | CUMV 27018 | 3.606 | -76.868 | KR817480 |
| 92 | Colombia, Valle del Cauca, Buenaventura, El Placer | CUMV 27020 | 3.606 | -76.868 | KR817516 |
| 93 | Colombia, Valle del Cauca, Buenaventura, El Placer | CUMV 27024 | 3.606 | -76.868 | KR817489 |
| 94 | Colombia, Valle del Cauca, Buenaventura, El Placer | CUMV 27022 | 3.606 | -76.868 | KR817518 |
| 95 | Colombia, Valle del Cauca, Buenaventura, El Placer | CUMV 27024 | 3.606 | -76.868 | KR817489 |
| 96 | Colombia, Valle del Cauca, Buenaventura, El Placer | CUMV 27026 | 3.606 | -76.868 | KR817519 |
| 97 | Colombia, Valle del Cauca, Buenaventura, El Placer | CUMV 27027 | 3.606 | -76.868 | KR817520 |
| 98 | Colombia, Valle del Cauca, Buenaventura, El Placer | CUMV 27028 | 3.606 | -76.868 | KR817481 |
| 99 | Colombia, Valle del Cauca, Buenaventura, El Placer | CUMV 27029 | 3.606 | -76.868 | KR817495 |
| 100 | Colombia, Valle del Cauca, Buenaventura, El Placer | CUMV 27030 | 3.606 | -76.868 | KR817521 |
| 101 | Colombia, Valle del Cauca, Buenaventura, El Placer | CUMV 27031 | 3.606 | -76.868 | KR817522 |
| 102 | Colombia, Valle del Cauca, Buenaventura, El Placer | CUMV 27032 | 3.606 | -76.868 | KR817523 |
| 103 | Colombia, Valle del Cauca, Buenaventura, El Placer | CUMV 27033 | 3.606 | -76.868 | KR817524 |
| 104 | Colombia, Valle del Cauca, Buenaventura, El Placer | CUMV 27034 | 3.606 | -76.868 | KR817525 |
| 105 | Colombia, Valle del Cauca, Buenaventura, El Placer | CUMV 27035 | 3.606 | -76.868 | KR817526 |
| 106 | Colombia, Valle del Cauca, Buenaventura, El Placer | CUMV 27036 | 3.606 | -76.868 | KR817527 |
| 107 | Colombia, Valle del Cauca, Dagua, Queremal | CUMV 27044 | 3.517 | -76.717 | KR817488 |
| 108 | Colombia, Valle del Cauca, Dagua, La Elsa | CUMV 27046 | 3.583 | -76.774 | KR817494 |
| 109 | Colombia, Valle del Cauca, Buenaventura, Río Anchicaya | CUMV 27040 | 3.614 | -76.905 | KR817530 |
| 110 | Colombia, Valle del Cauca, Dagua, La Elsa | CUMV 27049 | 3.583 | -76.774 | KR817532 |
| 111 | Colombia, Valle del Cauca, Dagua, La Elsa | CUMV 27050 | 3.583 | -76.774 | KR817533 |
| 112 | Colombia, Valle del Cauca, Dagua, La Elsa | CUMV 27051 | 3.583 | -76.774 | KR817534 |
| 113 | Colombia, Valle del Cauca, Dagua, La Elsa | CUMV 27052 | 3.583 | -76.774 | KR817535 |
| 114 | Colombia, Valle del Cauca, Dagua, La Elsa | CUMV 27053 | 3.583 | -76.774 | KR817536 |
| 115 | Colombia, Valle del Cauca, Dagua, La Elsa | CUMV 27055 | 3.583 | -76.774 | KR817537 |
| 116 | Colombia, Valle del Cauca, Dagua, La Elsa | CUMV 27056 | 3.583 | -76.774 | KR817538 |
| 117 | Colombia, Valle del Cauca, Dagua, La Elsa | CUMV 27057 | 3.583 | -76.774 | KR817539 |
| 118 | Colombia, Valle del Cauca, Dagua, La Elsa | CUMV 27058 | 3.583 | -76.774 | KR817482 |
| 119 | Colombia, Valle del Cauca, Dagua, La Elsa | CUMV 27059 | 3.583 | -76.774 | KR817540 |
| 120 | Colombia, Valle del Cauca, Dagua, La Elsa | CUMV 27060 | 3.583 | -76.774 | KR817541 |
| 121 | Colombia, Valle del Cauca, Dagua, Rio Blanco | CUMV 27068 | 3.600 | -76.817 | KR817545 |
| 122 | Colombia, Valle del Cauca, Dagua, Rio Blanco | CUMV 27074 | 3.600 | -76.817 | KR817548 |
| 123 | Colombia, Valle del Cauca, Dagua, Rio Blanco | CUMV 27076 | 3.600 | -76.817 | KR817490 |
| 124 | Colombia, Valle del Cauca, Dagua, Rio Blanco | CUMV 27078 | 3.600 | -76.817 | KR817549 |
| 125 | Colombia, Valle del Cauca, Dagua, Rio Blanco | CUMV 27079 | 3.600 | -76.817 | KR817550 |
| 126 | Colombia, Valle del Cauca, Dagua, Rio Blanco | CUMV 27082 | 3.600 | -76.817 | KR817551 |
| 127 | Colombia, Valle del Cauca, Dagua, Rio Blanco | CUMV 27083 | 3.600 | -76.817 | KR817552 |
| 128 | Colombia, Valle del Cauca, Dagua, Rio Blanco | CUMV 27084 | 3.600 | -76.817 | KR817553 |
| 129 | Colombia, Valle del Cauca, Dagua, Salado | CUMV 26992 | 3.567 | -76.717 | KR817558 |
| 130 | Colombia, Valle del Cauca, Dagua, Salado | CUMV 26993 | 3.567 | -76.717 | KR817502 |
| 131 | Colombia, Valle del Cauca, Dagua, Salado | CUMV 26994 | 3.567 | -76.717 | KR817557 |
| 132 | Colombia, Valle del Cauca, Dagua, Salado | CUMV 26995 | 3.567 | -76.717 | KR817492 |
| 133 | Colombia, Valle del Cauca, Dagua, Salado | CUMV 26996 | 3.567 | -76.717 | KR817503 |
| 134 | Colombia, Valle del Cauca, Dagua, Salado | CUMV 26997 | 3.567 | -76.717 | KR817497 |
| 135 | Colombia, Valle del Cauca, Dagua, Salado | CUMV 26998 | 3.567 | -76.717 | KR817504 |
| 136 | Colombia, Valle del Cauca, Dagua, Salado | CUMV 26999 | 3.567 | -76.717 | KR817505 |
| 137 | Colombia, Valle del Cauca, Dagua, Salado | CUMV 27000 | 3.567 | -76.717 | KR817506 |
| 138 | Colombia, Valle del Cauca, Dagua, Salado | CUMV 27001 | 3.567 | -76.717 | KR817507 |
| 139 | Colombia, Valle del Cauca, Dagua, Salado | CUMV 27002 | 3.567 | -76.717 | KR817484 |
| 140 | Colombia, Valle del Cauca, Dagua, Salado | CUMV 27003 | 3.567 | -76.717 | KR817508 |
| 141 | Colombia, Valle del Cauca, Dagua, Salado | CUMV 27004 | 3.567 | -76.717 | KR817509 |
| 142 | Colombia, Valle del Cauca, Dagua, Salado | CUMV 27005 | 3.567 | -76.717 | KR817510 |
| 143 | Colombia, Valle del Cauca, Dagua, Salado | CUMV 27006 | 3.567 | -76.717 | KR817493 |
| 144 | Colombia, Valle del Cauca, Dagua, Salado | CUMV 27007 | 3.567 | -76.717 | KR817485 |
| 145 | Colombia, Valle del Cauca, Dagua, Salado | CUMV 27008 | 3.567 | -76.717 | KR817511 |
| 146 | Colombia, Valle del Cauca, Dagua, Salado | CUMV 27010 | 3.567 | -76.717 | KR817487 |
| 147 | Colombia, Valle del Cauca, Dagua, Salado | CUMV 27011 | 3.567 | -76.717 | KR817512 |
| 148 | Colombia, Valle del Cauca, Dagua, Salado | CUMV 27012 | 3.567 | -76.717 | KR817486 |
| 149 | Colombia, Valle del Cauca, Dagua, Queremal | CUMV 27045 | 3.517 | -76.717 | KR817496 |
| 150 | Colombia, Valle del Cauca, La Cumbre, San Antonio | CUMV 22956 | 3.500 | -76.633 | KR817498 |
| 151 | Colombia, Valle del Cauca, La Cumbre, San Antonio | CUMV 22958 | 3.500 | -76.633 | KR817499 |
| 152 | Colombia, Valle del Cauca, Palmira, La Manuelita | CUMV 22959 | 3.583 | -76.283 | KR817500 |
| 153 | Colombia, Valle del Cauca, Palmira, Miraflores | CUMV 22961 | 3.578 | -76.434 | KR817501 |
| 154 | Colombia, Valle del Cauca, Tuluá, La Paila | CUMV 27086 | 4.319 | -76.072 | KR817554 |
| 155 | Colombia, Valle del Cauca, Jamundi, La Olga | CUMV 27087 | 3.299 | -76.519 | KR817555 |
| 156 | Colombia, Valle del Cauca, Buga, La Habana | CUMV 27088 | 3.881 | -76.191 | KR817556 |

## References

1. Barton NH, Hewitt GM. Analysis of hybrid zones. Annu Rev Ecol Syst 1985;16:113–48.

2. Coyne JA, Orr HA. Speciation. Sunderland: Sinauer Associates; 2004.

3. Durrett R, Buttel L, Harrison R. Spatial models for hybrid zones. Heredity. 2000;84:9–19.

4. Mayr E. Systematics and the origin of species, from the viewpoint of a zoologist. Harvard University Press; 1942.

5. Harrison RG. Hybrid zones: windows on evolutionary process. In: Futuyma DJ, Antonovics J, editors. Oxford surveys in evolutionary biology. Oxford: Oxford University Press; 1990. pp. 69–128.

6. Vines TH, Köhler SC, Thiel M, Ghira I, Sands TR, MacCallum CJ, et al. The maintenance of reproductive isolation in a mosaic hybrid zone between the fire-bellied toads *Bombina bombina and B.variegata*. Evolution. 2003;57:1876–88.

7. Endler JA. Geographic variation, speciation, and clines. Princeton University Press; 1977.

8. Peterson AT, Martínez-Meyer E, González-Salazar C. Reconstructing the Pleistocene geography of the *Aphelocoma* jays (Corvidae). Drivers Distrib. 2004;10:237–46.

9. Carstens BC, Richards CL. Integrating coalescent and ecological niche modeling in comparative phylogeography. Evolution. 2007;61:1439–54.

10. Richards CL, Carstens BC, Knowles L. Distribution modelling and statistical phylogeography: an integrative framework for generating and testing alternative biogeographical hypotheses. J Biogeog. 2007;34:1833–45.

11. Carnaval AC, Hickerson MJ, Haddad CFB, Rodrigues MT, Moritz C. Stability predicts genetic diversity in the Brazilian Atlantic forest hotspot. Science. 2009;323:785–9.

12. Swenson NG. Gis-based niche models reveal unifying climatic mechanisms that maintain the location of avian hybrid zones in a North American suture zone. J Evol Biol 2006;19:717–25.

13. Ruegg KC, Hijmans RJ, Moritz C. Climate change and the origin of migratory pathways in the Swainson’s thrush, *Catharus ustulatus*. J Biogeog. 2006;33:1172–82.

14. Swenson NG. The past and future influence of geographic information systems on hybrid zone, phylogeographic and speciation research. J Evol Biol. 2008;21:421–34.

15. Moritz C, Hoskin CJ, MacKenzie JB, Phillips BL, Tonione M, Silva N, et al. Identification and dynamics of a cryptic suture zone in tropical rainforest. Proc R Soc Lond B Biol Sci. 2009;276:1235–44.

16. Moore BR, Price JT. Nature of selection in the northern flicker hybrid zone and its implications for speciation theory. In: Harrison RG, editor. Hybrid zones and the evolutionary process. Oxford University Press; 1993. pp. 196–225.

17. Kruuk LEB, Baird SJE, Gale KS, Barton NH. A comparison of multilocus clines maintained by environmental adaptation or by selection against hybrids. Genetics. 1999;153:1959–71.

18. Buggs RJA. Empirical study of hybrid zone movement. Heredity. 2007;99:301–12.

19. Rohwer S, Bermingham E, Wood C. Plumage and mitochondrial DNA haplotype variation across a moving hybrid zone. Evolution. 2001;55:405–22.

20. Krosby M, Rohwer S. A 2000 km genetic wake yields evidence for northern glacial refugia and hybrid zone movement in a pair of songbirds. Proc R Soc Lond B Biol Sci. 2009;276:615–21.

21. Krosby M, Rohwer S. Ongoing movement of the Hermit warbler X Townsend’s warbler hybrid zone. PLoS One; 2010;5:e14164.

22. Taylor SA, Larson EL, Harrison RG. Hybrid zones: windows on climate change. Trends Ecol Evolut. 2015;30:398–406.

23. Leaché AD, Grummer JA, Harris RB, Breckheimer IK. Evidence for concerted movement of nuclear and mitochondrial clines in a lizard hybrid zone. Mol Ecol. 2017;23:75.

24. Griscom L. Notes on imaginary species of *Ramphocelus*. Auk. 1932;49:199–203.

25. Sibley CG. Hybridization in some colombian tanagers, avian genus *Ramphocelus*. Proc Am Philos Soc. 1958;102:448–53.

26. Olson SL, Violani C. Some unusual hybrids of *Ramphocelus*, with remarks on evolution in the genus (Aves: Thraupinae). Boll Mus Regionale Sci Nat Torino. 1995;13:297–312.

27. McCarthy EM. Handbook of avian hybrids of the world. Oxford University Press, USA; 2006.

28. Chapman FM. The distribution of bird-life in Colombia; a contribution to a biological survey of South America. Bull Am Mus Nat Hist. 1917;36:1–729.

29. Bedoya MJ, Murillo OE. Evidencia morfológica de hibridación entre las subespecies de Ramphocelus flammigerus (Passeriformes: Thraupidae) en Colombia. Rev Biol Trop. 2012;60:75–85.

30. Remsen JV Jr, Areta JI, Cadena CD, Claramunt S, Jaramillo A, Pacheco JF, et al. A classification of the bird species of South America. American Ornithologists’ Union. 2017: Available from: http://www.museum.lsu.edu/˜Remsen/SACCBaseline.html

31. Plath K. Color change in Ramphocelus flammigerus. Auk. 1945;62:304.

32. Brush AH. Pigments in hybrid, variant and melanic tanagers (birds). Comp Biochem Physiol. 1970;36:785–93.

33. Krueger TR, Williams DA, Searcy WA. The genetic mating system of a tropical tanager. Condor. 2008;110:559–62.

34. Keller LF, Grant PR, Grant BR, Petren K. Heritability of morphological traits in Darwin’s finches: misidentified paternity and maternal effects. Heredity. 2001;87:325–36.

35. Bears H, Drever MC, Martin K. Comparative morphology of Dark-eyed juncos Junco hyemalis breeding at two elevations: a common aviary experiment. J Avian Biol. 2008;39:152–62.

36. Ballentine B, Greenberg R. Common garden experiment reveals genetic control of phenotypic divergence between swamp sparrow subspecies that lack divergence in neutral genotypes. PLoS One. 2010;5:e10229.

37. Schluter D, Smith JNM. Natural selection on beak and body size in the song sparrow. Evolution. 1986;40:221–31.

38. Grant PR, Grant BR. Unpredictable evolution in a 30-year study of Darwin’s finches. Science. 2002;296:707–11.

39. Benkman CW. Divergent selection drives the adaptive radiation of crossbills. Evolution. 2003;57:1176–81.

40. Brumfield RT, Jernigan RW, McDonald DB, Braun MJ. Evolutionary implications of divergent clines in an avian (Manacus: Aves) hybrid zone. Evolution. 2001;55:2070–87.

41. Baldassarre DT, White TA, Karubian J, Webster MS. Genomic and morphological analysis of a semipermeable avian hybrid zone suggests asymmetrical introgression of a sexual signal. Evolution. 2014;68:2644–57.

42. Carling MD, Brumfield RT. Speciation in Passerina buntings: introgression patterns of sex-linked loci identify a candidate gene region for reproductive isolation. Mol Ecol. 2009;18:834–47.

43. Taylor SA, Curry RL, White TA, Ferretti V, Lovette IJ. Spatiotemporally consistent genomic signatures of reproductive isolation in a moving hybrid zone. Evolution. 2014;68:3066–81.

44. Gowen FC, Maley JM, Cicero C, Peterson AT, Faircloth BC, Warr TC, et al. Speciation in Western Scrub-Jays, Haldane’s rule, and genetic clines in secondary contact. BMC Evol Biol.2014; 14:135.

45. Darwin Database. BIOMAP Alliance Partners. 2007: Available from:http://www.biomap.net/BioMAP/login_en.php

46. Hijmans RJ, Cameron SE, Parra JL, Jones PG, Jarvis A. Very high resolution interpolated climate surfaces for global land areas. Int J Climatol. 2005;25:1965–78.

47. Phillips SJ, Anderson RP, Schapire RE. Maximum entropy modeling of species geographic distributions. Ecol Modell. 2006;190:231–59.

48. Fielding AH, Bell JF. A review of methods for the assessment of prediction errors in conservation presence/absence models. Environ Conserv. 1997;24:38–49.

49. Dinerstein E. A conservation assessment of the terrestrial ecoregions of Latin America and the Caribbean. Washington, D.C.: World Bank; 1995.

50. Hackett SJ. Molecular phylogenetics and biogeography of tanagers in the genus Ramphocelus (Aves). Mol Phylogenet Evol. 1996;5:368–82.

51. Burns KJ, Racicot RA. Molecular phylogenetics of a clade of lowland tanagers: implications for avian participation in the Great American Interchange. Auk. 2009;126:635–48.

52. Sambrook J, Russell DW. Molecular cloning: a laboratory manual. Third Edition. Cold Spring Harbor, N.Y.: Cold Spring Harbor Laboratory; 2001.

53. Sorenson MD, Ast JC, Dimcheff DE, Yuri T, Mindell DP. Primers for a PCR-based approach to mitochondrial genome sequencing in birds and other vertebrates. Mol Phylogenet and Evol.1999;12:105–14.

54. Carling MD, Serene LG, Lovette IJ. Using historical DNA to characterize hybridization between Baltimore Orioles (Icterus galbula) and Bullock’s Orioles (I. bullockii). Auk. 2011;128:61–8.

55. Stamatakis A, Hoover P, Rougemont J. A rapid bootstrap algorithm for the RAxML Web servers.Syst Biol. 2008;57:758–71.

56. Sato A, Tichy H, O’hUigin C, Grant PR, Grant BR, Klein J. On the origin of Darwin’s finches. Mol Biol Evol. 2001;18:299–311.

57. Excoffier L, Lischer HEL. Arlequin suite ver 3.5: a new series of programs to perform population genetics analyses under Linux and Windows. Mol Ecol Resour.2010;10:564–7.

58. Heled J, Drummond AJ. Bayesian inference of population size history from multiple loci. BMC Evol Biol. 2008;8:289.

59. Darriba D, Taboada GL, Doallo R, Posada D. jModelTest 2: more models, new heuristics and parallel computing. Nat Methods. 2012;9:772.

60. Weir JT, Price M. Andean uplift promotes lowland speciation through vicariance and dispersal in Dendrocincla woodcreepers. Mol Ecol. 2011;20:4550–63.

61. R Development Core Team. R: A language and environment for statistical computing. R Foundation for Statistical Computing, Vienna, Austria. http://www.R-project.org/. 2013.

62. Valderrama E, Pérez-Emán JL, Brumfield RT, Cuervo AM, Cadena CD. The influence of the complex topography and dynamic history of the montane Neotropics on the evolutionary differentiation of a cloud forest bird (Premnoplex brunnescens, Furnariidae). J Biogeog. 2014;41:1533–46.

63. Endler JA. On the measurement and classification of colour in studies of animal colour patterns. Biol J Linn Soc. 1990;41:315–52.

64. Parra JL. Color evolution in the hummingbird genus Coeligena. Evolution. 2009;64:324–35.

65. Derryberry EP, Derryberry GE, Maley JM, Brumfield RT. HZAR: hybrid zone analysis using an R software package. Mol Ecol Resour. 2014;14:652–63.

66. Barton NH, Baird S. Analyse: software for the analysis of geographic variation and hybrid zones. University of Edinburgh; 1999.

67. Porter AH, Wenger R, Geiger H, Scholl A, Shapiro AM. The Pontia daplidice-edusa hybrid zone in northwestern Italy. Evolution. 1997;51:1561–73.

68. Ritz C, Baty F, Streibig JC, Gerhard D. Dose-Response Analysis Using R. PLoS One. 2015;10:e0146021.

69. Ritz C, Streibig JC. Bioassay analysis using R. J Stat Softw. 2005;12:1–22.

70. Hewitt G. The genetic legacy of the Quaternary ice ages. Nature. 2000;405:907–13.

71. Haffer J. Speciation in colombian forest birds west of the Andes. Am Mus Novit. 1967;2294:1–57.

72. Endler JA. Problems in distinguishing historical from ecological factors in biogeography. Am Zool. 1982;22:441–52.

73. Brumfield RT. Mitochondrial variation in bolivian populations of the Variable Antshrike (Thamnophilus caerulescens). Auk. 2005;122:414–32.

74. Dasmahapatra KK, Lamas G, Simpson F, MalletJ. The anatomy of a “suture zone” in Amazonian butterflies: a coalescent-based test for vicariant geographic divergence and speciation. Mol Ecol. 2010;19:4283–301.

75. Hooghiemstra H, Van der Hammen T. Quaternary Ice-Age dynamics in the Colombian Andes: developing an understanding of our legacy. Philos Trans R Soc Lond Biol Sci. 2004;359:173–81.

76. Ramírez-Barahona S, Eguiarte LE. The role of glacial cycles in promoting genetic diversity in the Neotropics: the case of cloud forests during the Last Glacial Maximum. Ecol Evol. 2013;3:725–38.

77. Cadena CD, Pedraza CA, Brumfield RT. Climate, habitat associations and the potential distributions of Neotropical birds: Implications for diversification across the Andes. Rev Acad Colomb Cienc Ex Fis Nat 2016;40:275.

78. Jansson R, Dynesius M. The fate of clades in a world of recurrent climatic change: Milankovitch oscillations and evolution. Annu Rev Ecol Syst 2002;33:741–77.

79. Arias CF, Rosales C, Salazar C, Castaño J, Bermingham E, Linares M, et al. Sharp genetic discontinuity across a unimodal Heliconius hybrid zone. Mol Ecol. 2012;21:5778–94.

80. Arias CF, Muñoz AG, Jiggins CD, Mavárez J, Bermingham E, Linares M. A hybrid zone provides evidence for incipient ecological speciation in Heliconius butterflies. Mol Ecol. 2008;17:4699–712.

81. Medina I, Wang IJ, Salazar C, Amézquita A. Hybridization promotes color polymorphism in the aposematic harlequin poison frog, Oophaga histrionica. Ecol Evol. 2013;3:4388–400.

82. Barton NH, Gale KS. Genetic analysis of hybrid zones. In: Harrison RG, editor. Hybrid zones and the evolutionary process. New York: Oxford Univ. Press; 1993. pp. 13–45.

83. Gay L, Crochet P-A, Bell DA, Lenormand T. Comparing clines on molecular and phenotypic traits in hybrid zones: a window on tension zone models. Evolution. 2008;62:2789–806.

84. Dasmahapatra KK, Blum MJ, Aiello A, Hackwel S, Davies N, Bermingham EP, et al. Inferences from a rapidly moving hybrid zone. Evolution. 2002;56:741–53.

85. Smith KL, Hale JM, Gay L, Kearney M, Austin JJ, Parris KM, et al. Spatio-temporal changes in the structure of an Australian frog hybrid zone: a 40-year perspective. Evolution. 2013;67:3442–54.

86. Mettler RD, Spellman GM. A hybrid zone revisited: molecular and morphological analysis of the maintenance, movement, and evolution of a Great Plains avian (Cardinalidae: *Pheucticus*) hybrid zone. Mol Ecol. 2009;18:3256–67.

87. Rosser N, Dasmahapatra KK, Mallet J. Stable *Heliconius* butterfly hybrid zones are correlated with a local rainfall peak at the edge of the Amazon basin. Evolution. 2014;68:3470–84.

88. McDonald DB, Clay RP, Brumfield RT, Braun MJ. Sexual selection on plumage and behavior in an avian hybrid zone: experimental tests of male-male interactions. Evolution. 2001;55:1443–51.

89. Bronson CL, Grubb TCJ, Sattler DG, Braun MJ. Mate preference: a possible causal mechanism for a moving hybrid zone. Anim Behav. 2003;65:489–500.

90. Stein AC, Uy JAC. Unidirectional introgression of a sexually selected trait across an avian hybrid zone: a role for female choice? Evolution. 2006;60:1476–85.

91. Toews DPL, Hofmeister NR, Taylor SA. The evolution and genetics of carotenoid processing in animals. Trends Genet. 2017;33:171–82.

92. Blount JD, McGraw KJ. Signal functions of carotenoid coloration. In: Britton G, Liaaen-Jensen S, editors. Carotenoids. Basel: Birkhauser; 2008. pp. 213–36.

93. Hey J. Multilocus methods for estimating population sizes, migration rates and divergence time, with applications to the divergence of *Drosophila pseudoobscura* and *D. persimilis*. Genetics. 2004;167:747–60.

94. Jankowski JE, Londoño GA, Robinson SK, Chappell MA. Exploring the role of physiology and biotic interactions in determining elevational ranges of tropical animals. Ecography. 2012;36:1–12.

95. Engler JO, Rödder D, Elle O, Hochkirch A, Secondi J. Species distribution models contribute to determine the effect of climate and interspecific interactions in moving hybrid zones. J Evol Biol. 2013;26:2487–96.

96. Chunco AJ. Hybridization in a warmer world. Ecol Evol. 2014;4:2019–31.

97. Billerman SM, Murphy MA, Carling MD. Changing climate mediates sapsucker (Aves: Sphyrapicus) hybrid zone movement. Ecol Evol. 2016;6:7976–90.

98. Taylor SA, White TA, Hochachka WM, Ferretti V, Curry RL, Lovette IJ. Climate-mediated movement of an avian hybrid zone. Curr Biol. 2014;24:671–6.

99. Parsons TJ, Olson SL, Braun MJ. Unidirectional spread of secondary sexual plumage traits across an avian hybrid zone. Science. 1993;260:1643–6.

100. Ruegg KC. Genetic, morphological, and ecological characterization of a hybrid zone that spans a migratory divide. Evolution. 2008;62:452–66.

101. Carling MD, Zuckerberg B. Spatio-temporal changes in the genetic structure of the *Passerina* bunting hybrid zone. Mol Ecol. 2011;20:1166–75.

102. Milá B, Toews DPL, Smith BT, Wayne RK. A cryptic contact zone between divergent mitochondrial DNA lineages in southwestern North America supports past introgressive hybridization in the yellow-rumped warbler complex (Aves: *Dendroica coronata*). Biol J Linn Soc. 2011;103:696–706.

103. Miller MJ, Lipshutz SE, Smith NG, Bermingham E. Genetic and phenotypic characterization of a hybrid zone between polyandrous Northern and Wattled Jacanas in Western Panama. BMC Evol Biol.2014;14:227.

104. Engebretsen KN, Barrow LN, Rittmeyer EN, Brown JM, Moriarty Lemmon E. Quantifying the spatiotemporal dynamics in a chorus frog (Pseudacris) hybrid zone over 30 years. Ecol Evol.2016;6:5013–31.

105. Poelstra JW, Vijay N, Bossu CM, Lantz H, Ryll B, Müller I, et al. The genomic landscape underlying phenotypic integrity in the face of gene flow in crows. Science. 2014;344:1410–4.

106. Toews DPL, Taylor SA, Vallender R, Brelsford A, Butcher BG, Messer PW, et al. Plumage genes and little else distinguish the genomes of hybridizing warblers. Curr. Biol. 2016;26:2313–8.

